# Tumor cell-derived spermidine promotes a pro-tumorigenic immune microenvironment in glioblastoma via CD8+ T cell inhibition

**DOI:** 10.1101/2023.11.14.567048

**Authors:** Kristen E. Kay, Juyeun Lee, Ellen S. Hong, Julia Beilis, Sahil Dayal, Emily Wesley, Sofia Mitchell, Sabrina Z. Wang, Daniel J. Silver, Josephine Volovetz, Sadie Johnson, Mary McGraw, Matthew M. Grabowski, Tianyao Lu, Lutz Freytag, Vinod Narayana, Saskia Freytag, Sarah A. Best, James R. Whittle, Zeneng Wang, Ofer Reizes, Jennifer S. Yu, Stanley L. Hazen, J. Mark Brown, Defne Bayik, Justin D. Lathia

**Author notes:** **Corresponding author,** Justin D. Lathia 9500 Euclid Ave, NE3-202 Cleveland, OH 44195. **Disclosures:** The authors declare no competing interests.

## Abstract

The glioblastoma microenvironment is enriched in immunosuppressive factors that potently interfere with the function of cytotoxic T lymphocytes. Cancer cells can directly impact the immune system, but the mechanisms driving these interactions are not completely clear. Here we demonstrate that the polyamine metabolite spermidine is elevated in the glioblastoma tumor microenvironment. Exogenous administration of spermidine drives tumor aggressiveness in an immune-dependent manner in pre-clinical mouse models via reduction of CD8+ T cell frequency and phenotype. Knockdown of ornithine decarboxylase, the rate-limiting enzyme in spermidine synthesis, did not impact cancer cell growth in vitro but did result in extended survival. Furthermore, glioblastoma patients with a more favorable outcome had a significant reduction in spermidine compared to patients with a poor prognosis. Our results demonstrate that spermidine functions as a cancer cell-derived metabolite that drives tumor progression by reducing CD8+T cell number and function.

## Introduction

Despite aggressive multimodal therapies including maximal safe surgical resection followed by concomitant radiation and chemotherapy, patients with glioblastoma (GBM), the most common primary malignant brain tumor, continue to have a poor prognosis^1–3^. While advances, including targeted therapies and more recently immunotherapy, have been achieved in other advanced cancers, GBM outcomes have not changed dramatically in decades^4–6^. GBM remains a major clinical challenge due to a variety of unique barriers, including inherent tumor cell therapeutic resistance, an immune-suppressive microenvironment, and metabolic adaptability^7–10^. In particular, the tumor microenvironment contains elevated numbers of immunosuppressive cells and a limited amount of effector cells^11,12^. Moreover, tumor cells leverage bi-directional communication mechanisms to alter the immune microenvironment^13,14^. A better understanding of these communication mechanisms in the context of immune cell infiltration, as well as their impact on the balance between immune activation and suppression, is critical for a better understanding not only of the immune microenvironment but also of the tumor’s response within.

Metabolic alterations are a hallmark of cancer and are well characterized in GBM cells. Such changes include specific dependencies involving glycolysis and lipid metabolism^15,16^. Recent studies have demonstrated that metabolic programs are not static but are subject to plasticity and underlie cellular states that drive tumor growth and therapeutic resistance^17^. Metabolic alterations extend beyond tumor cells and impact immune cells as well, altering their function^18^. These immune cell-specific metabolic changes are triggered by the tumor microenvironment as well as tumor cells, representing another important cell communication mechanism that can alter tumor growth^19^.

Polyamines are a family of cationic metabolites that include putrescine, spermine, and spermidine. These metabolites can be generated from arginine and are produced by nearly every cell in the body. Polyamines are critical to many cellular homeostatic functions, including cell growth and proliferation through their role in DNA replication and translation^20^. In many cancers, including GBM, polyamines are elevated and support cancer cell growth and immune suppression^21^. Specifically, in GBM, it has been shown that the polyamine family member spermidine (SPD) increases the acidity of the tumor microenvironment, shifting the balance towards immune-suppressive myeloid cells^22^. Targeting the polyamine pathway at the rate-limiting step in biosynthesis has been demonstrated to increase survival in pre-clinical models of neuroblastoma and to synergize with conventional immune checkpoint inhibitor-based immunotherapies^23^. In pediatric gliomas, additional pre-clinical benefit was observed using a polyamine transport inhibitor in conjunction with biosynthesis disruption^24^. While these and other studies have demonstrated elevation of polyamines in GBM and a function in brain tumors, mainly involving myeloid cells, the specific sources of polyamines and the impact on the immune system as a whole are less clear. Here we show that increased SPD in the tumor microenvironment, produced in part from cancer cells, drives tumor progression by decreasing CD8+ T cell frequency and activity via reduced cytotoxic ability and increased apoptosis-induced death of CD8+ T cells.

## Results

### Spermidine drives GBM progression

It has previously been reported that GBM patients have increased SPD in their cerebrospinal fluid (CSF) and blood compared to healthy controls^25^. To investigate the extent to which this is paralleled in our pre-clinical mouse models, we intracranially implanted the mouse glioma models SB28 and GL261 into wild-type C57BL/6 mice. Mass spectrometry of tumor tissue from these mice revealed an increase in members of the polyamine family, including a substantial increase in SPD in the tumor setting compared to control conditions. We also observed a higher magnitude elevation in SPD in the brain compared to other polyamine family members (**Fig. 1A-D**). Furthermore, spatial MALDI-TOF analysis of an independent CT-2A glioma model revealed tumor-intrinsic production of SPD (**Supp Fig. S1A-E**) and related enzymes (**Supp Fig. S2A-G**). These findings corroborate previous observations in human patients and suggest that SPD is increased in the tumor microenvironment. Based on the sex differences observed in GBM, both epidemiologically and in terms of immune responses^26,27^, we assessed equal numbers of male and female mice and did not observe any substantial sex differences (**Supp Fig. S3A-E**). Given the lack of sex differences in response to SPD and the higher incidence and poorer outcome of GBM in males^28^, we focused on males for the subsequent studies. In order to explore what effect elevated SPD would have on tumor growth, we developed an experimental paradigm in which we intracranially implanted mouse glioma cells, as previously described, and administered SPD at regular intervals via intraperitoneal injection (**Fig. 1E**). We confirmed via mass spectrometry that mice receiving systemic SPD treatment had an increase in SPD levels within the tumor microenvironment, recapitulating a high SPD-producing tumor (**Fig. 1F**). Additionally, systemic endogenous treatment with SPD robustly shortened survival of immune-competent mice (**Fig. 1G-H**, **Supp Fig. S4A-D).** Taken together, these data suggest that SPD is elevated in the GBM microenvironment and accelerates GBM progression.

**Figure 1.**
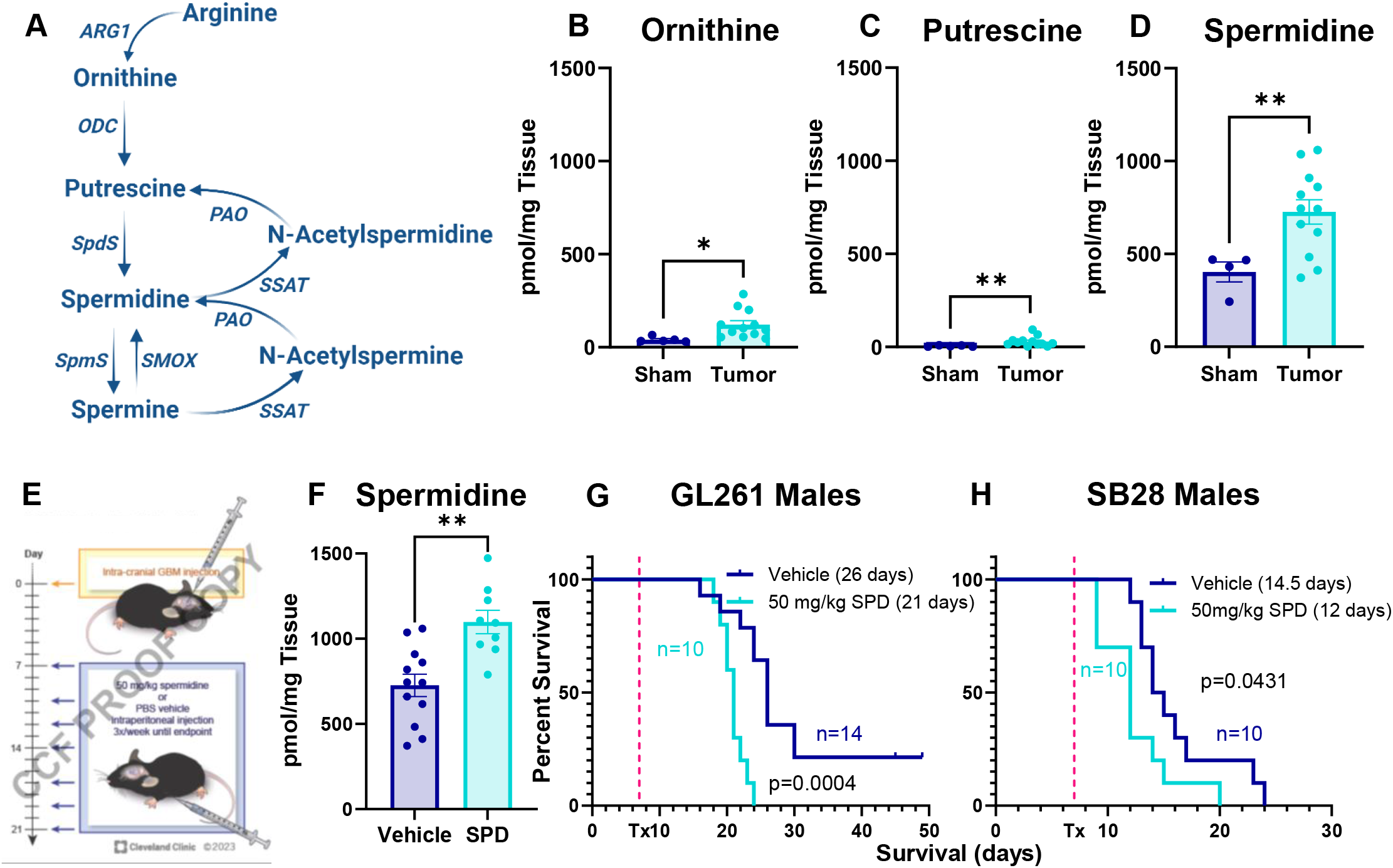
SPD levels are increased in mouse GBM models and drive GBM progression. (A) Polyamine biosynthesis pathway. (B-D) LC-MS was performed on tumors removed from B6 mice 17 days after intracranial injection of mouse GBM cell lines (25K/injection GL261). (E) Experimental paradigm for subsequent mouse experiments receiving tumor implantation followed by 50 mg/kg SPD IP treatment or PBS vehicle. (F) LC-MS/MS of tumor-bearing mice treated with IP SPD. (G-H) Survival analysis was performed after intracranial injection of mouse GBM cell lines (25K/injection GL261, 20K/injection SB28) in B6 mice. Median survival days and number of animals are indicated in the graph. Data combined from three independent experiments. Statistical significance for (B-D), (F) was determined by unpaired *t-*test (\**p*<0.05, \*\**p*<0.01). Statistical significance for (G-H) was determined by log-rank test, considering *p*-value <0.05 to be significant. ARG1: arginase, ODC: ornithine decarboxylase, SpdS: spermidine synthase, SpmS: spermine synthase, SSAT: spermidine/ spermine acetyl transferase, PAO: polyamine oxidase.

### Spermidine drives GBM growth in an immune-dependent manner

As SPD is involved in many cellular functions, including cell growth, we tested whether SPD has a direct effect on cancer cell growth. When mouse glioma cells were cultured in vitro with SPD for 72 hours, we observed no significant changes in cell numbers compared to control treatment (**Fig. 2A-B**). Additionally, the proliferation rate of brain resident populations (astrocytes, microglia) as well as human GBM and prostate cancer cells was not affected by the addition of exogenous SPD focus to other components of the tumor microenvironment that could contribute to the observed survival phenotype. GBM creates an immune-suppressive microenvironment characterized by an increase in immune-suppressive myeloid cells and limited T and NK cell infiltration^29,30^. Moreover, polyamines were recently shown to be critical for myeloid-driven immune suppression in GBM and T cell differentiation^11,22,31^. To investigate whether SPD could be altering immune cells, we repeated the same in vivo experimental paradigm previously described (**Fig. 1E**) using immunocompromised NSG (NOD.Cg-PrkdcscidIl2rgtm1Wjl/SzJ) mice. The sharp decline in survival observed in immune-competent mice with SPD treatment was abrogated in NSG mice, indicating that increased SPD is likely interfering with the immune response (**Fig. 2C-D, Supp Fig. 6A-D**). These data suggest that SPD likely drives tumor growth in an immune cell-dependent manner.

**Figure 2.**
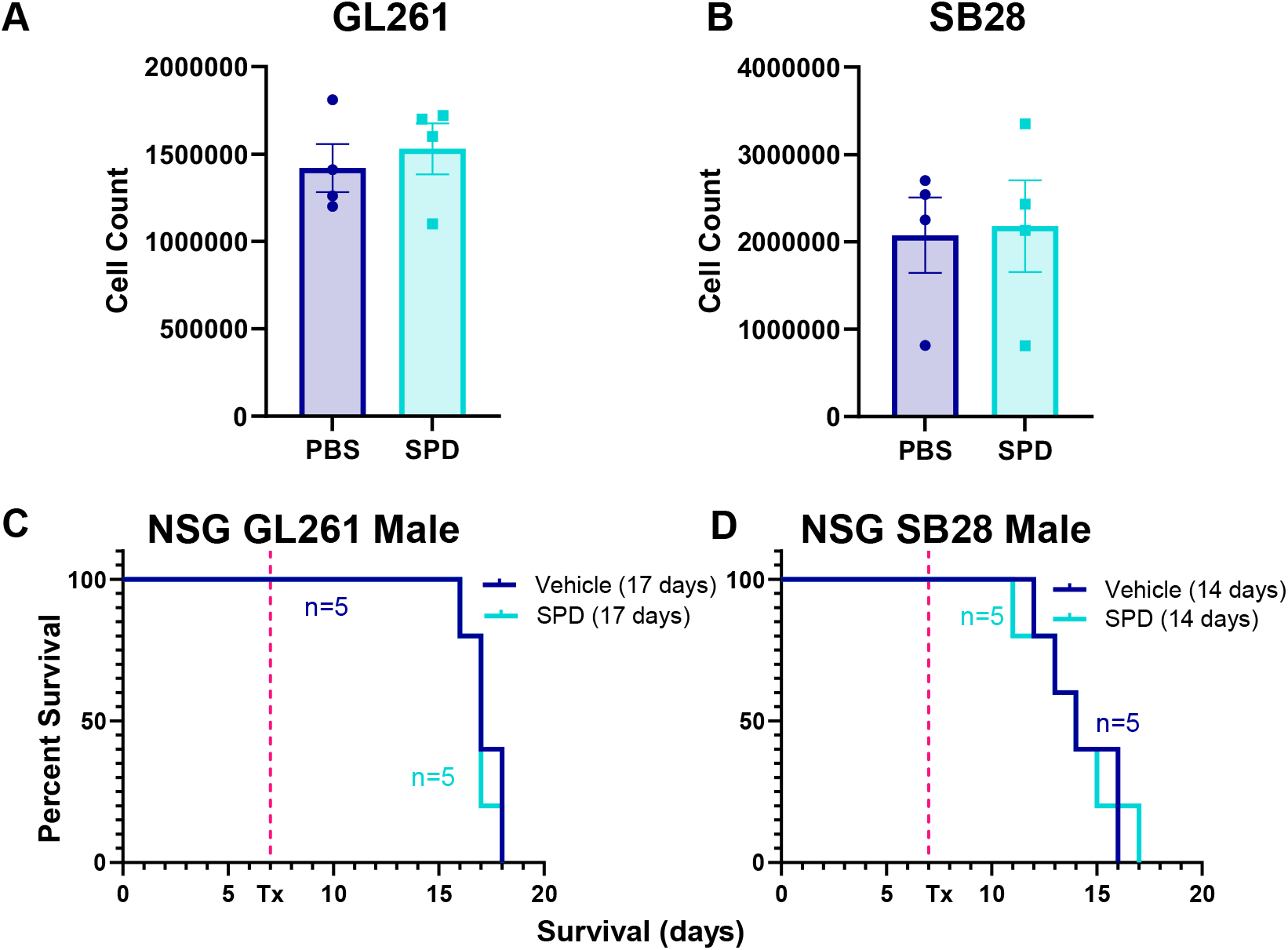
SPD interacts with the immune system to drive GBM progression. (A-B) Mouse glioma cells treated with physiological levels of SPD *in vitro* for 72 hours; data representative of 3 independent experiments. (C-D) Survival analysis was performed after intracranial injection of mouse GBM cell lines (25K/injection GL261, 20K/injection SB28) in immunocompromised male NSG mice, followed by 50 mg/kg SPD IP treatment or PBS vehicle. Median survival days and number of animals are indicated in the graph. Statistical significance was determined by log-rank test, considering *p*-value <0.05 to be significant.

### Spermidine drives GBM growth by reducing T cells

Based on previous reports indicating that SPD drives CD4+ T cell differentiation^31^, we investigated the effect of SPD on adaptive immune cells. Mouse splenocyte-derived lymphocytes treated with SPD in vitro and sorted via flow cytometry showed a significant reduction in both viable CD8+ and CD4+ T cells (**Fig. 3A-B**), as well as in B cells and NK cells (**Supp Fig. 7A-C**). To determine whether lymphocytes were driving SPD-mediated accelerated GBM growth in our mouse models, we repeated the same experimental paradigm (**Fig. 1E**) as described above in Rag1 knockout mice, which lack functional B and T cells. We observed no difference in survival between SPD and control treatment groups, supporting the hypothesis that SPD interacts with these immune cell subsets to drive GBM progression (**Fig. 3C-D, Supp Fig. 7D-G**).

**Figure 3.**
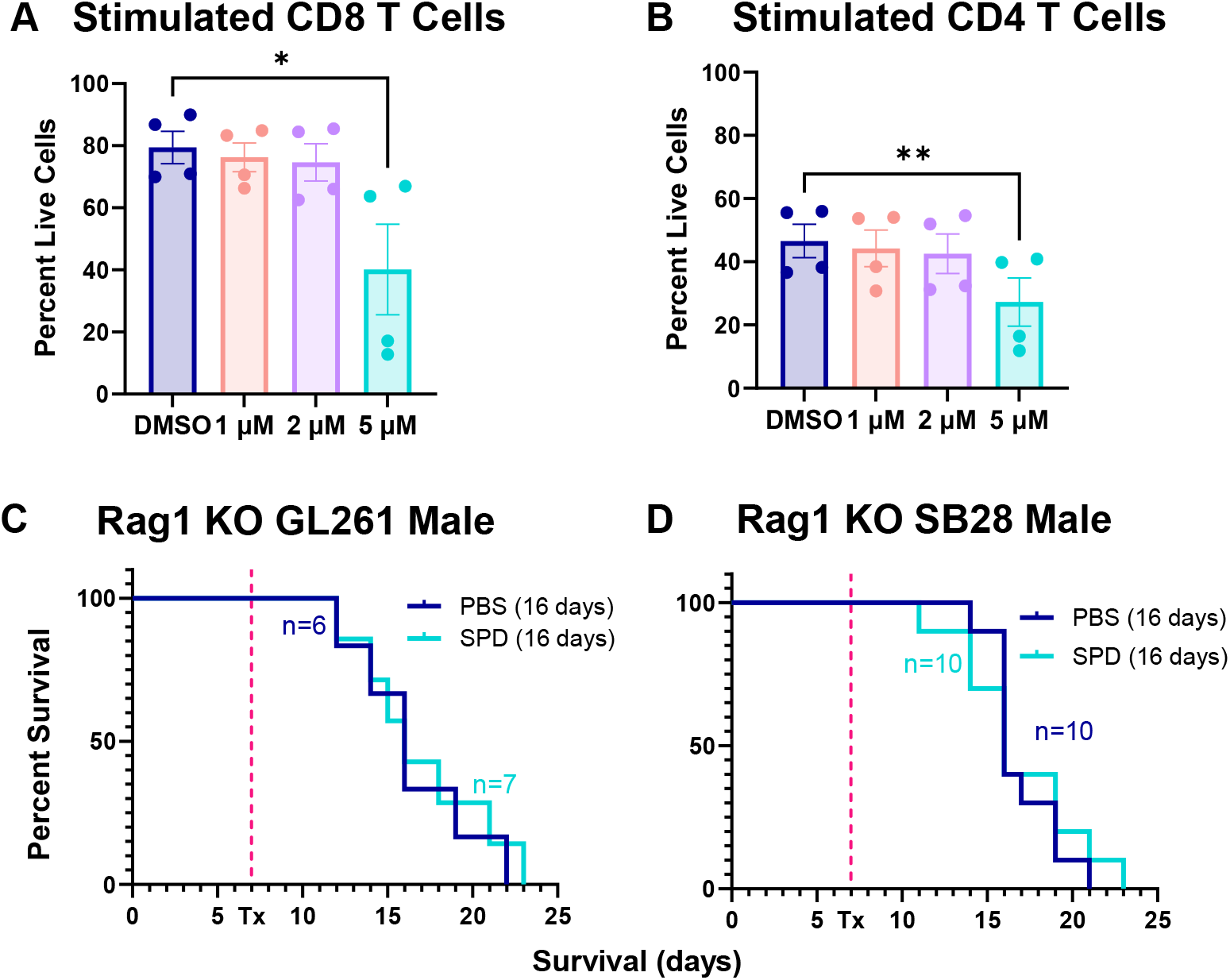
Lymphocyte subsets are affected by SPD. (A-B) Splenocyte-derived lymphocyte subsets were treated with physiological levels of SPD in vitro. (C-D) Survival analysis was performed after intracranial injection of mouse GBM cell lines (25K/injection GL261, 20K/injection SB28) in male Rag1 knockout mice, followed by 50 mg/kg SPD IP treatment or PBS vehicle. Median survival days and number of animals are indicated in the graph. Data combined from two independent experiments. Statistical significance for (A-B) was determined by one-way ANOVA (*p<0.05, **p<0.01). Statistical significance for (C-D) was determined by log-rank test, considering p-value <0.05 to be significant.

We then investigated changes to the immune response in the GBM microenvironment of immune-competent mice treated with exogenous SPD compared to control conditions. In the tumor-bearing hemisphere, we observed a significant reduction in the CD8/T regulatory cell (Treg) ratio, indicating decreased cytotoxic immune response in SPD-treated mice (**Fig. 4A**). This is partially due to the increased proportion of Tregs and a trend of decreasing of CD8+ T cell abundance (**Fig. 4B-C**). Additionally, we observed increased exhaustion markers specifically on CD8+ T cells in SPD-treated mice (**Fig. 4 D-E**). Immune analysis of blood and bone marrow replicated the immunosuppressive phenotype seen in the tumor tissue (**Supp Fig. S8A-I**). Treg exhaustion markers were not affected by SPD treatment (**Supp Fig. S8J-M**). Immune phenotyping of tumor-bearing mice suggests that increased SPD levels in the tumor microenvironment affect CD8+ T cells and Tregs, contributing to GBM progression (representative gating strategy in **Supp Fig. 9**).

**Figure 4.**
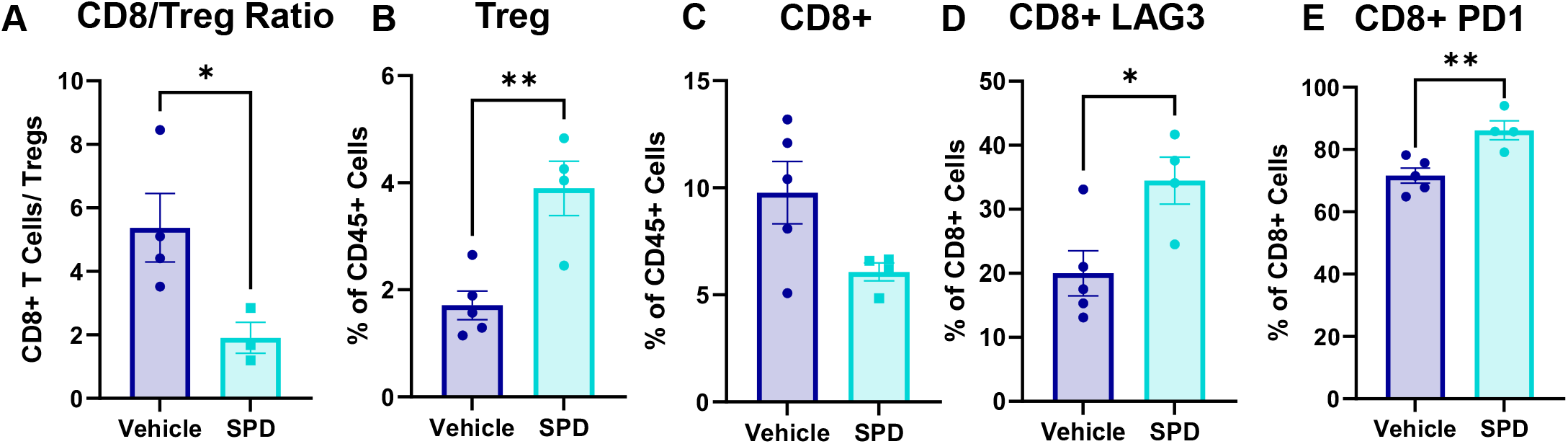
Exogenous treatment with SPD decreases cytotoxicity of CD8+ T cells. After intracranial injection of mouse GBM cell line SB28 (20K/injection) into male B6 mice followed by 50 mg/kg SPD IP treatment or PBS vehicle, the tumor-bearing hemisphere was collected and processed for flow cytometry immune phenotyping. (A) Ratio of CD8+ T cells and CD4+ Tregs. (B-C) Proportion of T cells in CD45+ cells. (D-E) Exhaustion markers of CD8+ T cells Statistical significance for (A-E) was determined by unpaired *t-* test (\**p*<0.05, \*\**p*<0.01).

Taken together, these data demonstrate that SPD reduces cytotoxic T cell number and phenotype.

### Ornithine decarboxylase drives GBM cell-mediated tumor growth and T cell alterations

Given that exogenously administered SPD drives tumor growth and alters T cell number and phenotype, we wanted to assess how this functions in a GBM cell-intrinsic manner. Using shRNA lentiviral particles targeting *ODC1*, the gene that encodes ornithine decarboxylase (ODC) – the rate-limiting irreversible enzyme of the main polyamine biosynthesis pathway – we knocked down *ODC1* in SB28 tumor cells (**Fig. 5A**), which resulted in decreased SPD production (**Fig. 5B**) and no significant changes in intrinsic tumor cell growth (**Fig. 5C**). Intracranial implantation of *ODC1*-knockdown GBM cells resulted in significantly extended survival compared to a non-target control (**Fig. 5D**), indicating that SPD production by cancer cells is partially responsible for GBM growth. Immune phenotyping of mice implanted with *ODC1*-knockdown cells revealed an increase in the proportion of CD8+ T cells in the TME compared to non-targeting controls (**Fig. 5E**). Additionally, CD8+ T cell exhaustion markers were decreased, suggesting the CD8+ T cells might have increased cytotoxicity in the tumor microenvironment (**Fig. 5F-G**). These data suggest that SPD generated by GBM cells via ODC can drive GBM growth and attenuate T cell number and function, which is consistent with our findings observed with exogenous administration of SPD.

**Figure 5.**
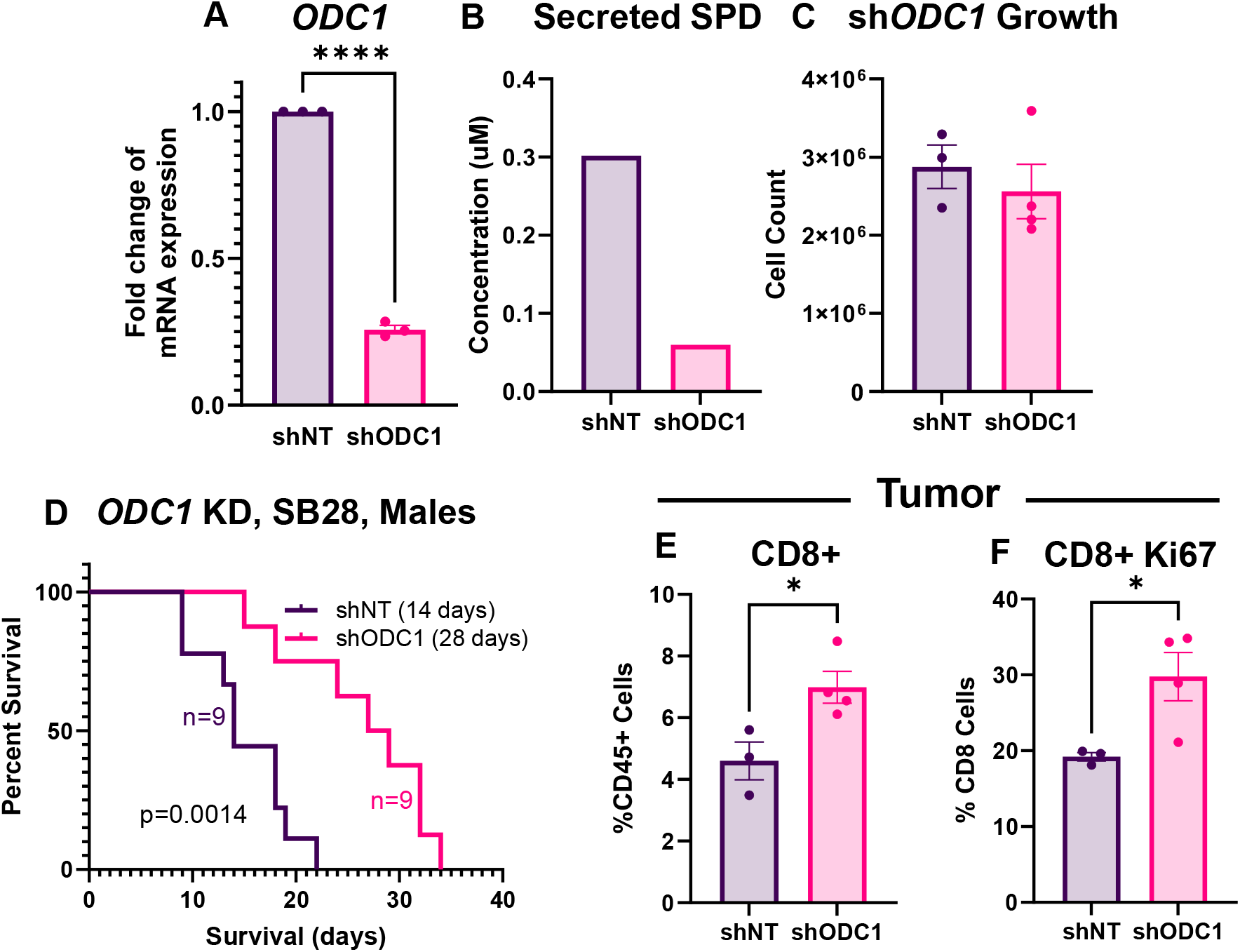
Knockdown of the polyamine biosynthesis pathway extends survival. (A) mRNA expression of *ODC1* in shRNA knockdown mouse glioma cells compared to non-targeted control. (B) Conditioned media SPD measurement via mass spectrometry. (C) Cell count after 72 hours growth. (D) Survival analysis was performed after intracranial injection of shRNA-modified mouse GBM cells (20K nontarget or *ODC1* KD SB28 cells) in B6 mice. Median survival days and number of animals are indicated in the graph. Data combined from two independent experiments. (E-F) Immune phenotyping via flow cytometry was performed on tumors removed from B6 mice 14 days after intracranial injection of shRNA-modified mouse GBM cells (20K nontarget or *ODC1* KD SB28 cells). (E) Percentage of CD8+ cells in tumor. (F) Proliferation marker in CD8+ T cells. Statistical significance for (D) was determined by log-rank test, considering *p*-value <0.05 to be significant. Statistical significance for (A, C, E-F) was determined by unpaired *t-*test (\**p*<0.05, ****p<0.0001).

### SPD induces CD8+ T cell apoptosis

To elucidate the mechanism through which SPD affects CD8+ T cells, we first investigated cell death and apoptosis, as SPD is known to be involved in apoptotic pathways^32^. Treating splenocyte-derived CD8+ T cells with SPD during the in vitro stimulation process for 72 hours resulted in an increase in fully apoptotic cells and a reduction in live cells (**Fig. 6A-C**). Additionally, no difference was noted in cell proliferation of CD8+ T cells treated with SPD compared with vehicle-treated cells (**Supp Fig. S10A**). CD8+ T cells treated in the same manner were analyzed for cytokine profile changes; we observed an increase in the exhaustion marker TIM3 as well as a reduction of the activation marker CD44 (**Fig. 6D-E**). The number of CD8+ T cells producing the established anti-tumorigenic cytokines IFNγ and TNFα was reduced (**Fig. 6F-G**). Taken together, these data suggest that SPD increases apoptosis, thus decreasing the available cytotoxic cells in the CD8+ T cell pool, in addition to decreasing their functionality by altering their cytokine profile to a less cytotoxic phenotype.

### SPD is correlated with decreased CD8+ T cells and a poorer prognosis

To investigate parallels between GBM patients and our preclinical findings, we interrogated multiple components of the SPD pathway and the tumor microenvironment. TCGA and GTEX data of normal brain tissue compared with low-grade glioma showed an increase in *ODC1* mRNA expression; when compared to GBM patients, there was a robust increase in expression in GBM compared to all other groups (**Fig. 7A**). To assess whether *ODC1* expression is linked to changes in the immune microenvironment, we analyzed single-cell RNAseq data from Ruiz-Moreno et al.^33^ and found that higher expression of *ODC1* in cancer cells correlated with fewer CD8+ T cells in the tumor microenvironment in GBM patients (**Fig. 7B**), similar to what we observed in mouse models. Furthermore, Visium spatial analysis of GBM patients from Ravi et al.^34^ showed a negative correlation between SPD-producing enzymes and the areas immediately surrounding identified CD8+ T cells (**Fig. 7C**). Finally, to link spermidine levels to GBM patient survival, tumor samples from age-matched GBM patients with shorter overall survival (median = 9.4 months) and longer-term overall survival (median = 41.4 months) were compared. We found significantly lower levels of spermidine present in the tumor microenvironment of GBM patients with longer survival outcomes (**Fig. 7D, Supp Fig. S11A-C**). Taken together, these data further reinforce that SPD is associated with poor GBM patient outcome and a reduction in CD8+ T cells in the tumor microenvironment.

**Figure 6.**
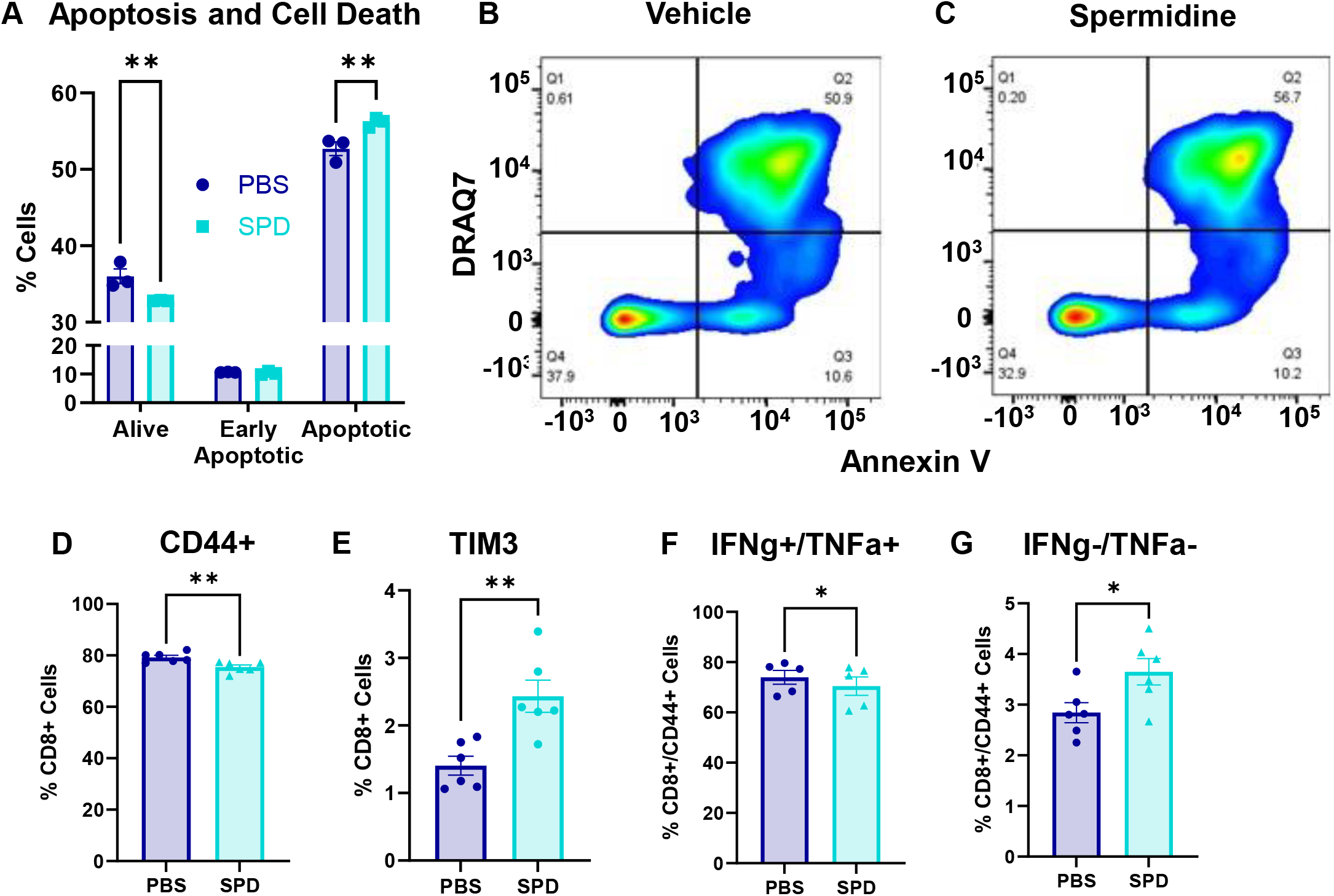
CD8+ T cells have reduced viability and functionality in the presence of SPD. Splenocyte-derived CD8+ T cells were treated with SPD *in vitro*. (A-C) Apoptotic cells and cell death were measured via Annexin V and DRAQ7 staining, respectively, and analyzed via flow cytometry. Statistical significance was determined by two-way ANOVA (\*\**p*<0.01). (B-C) Visual representation of gain in double-positive cells under SPD treatment. (D-E) T cell markers in CD8+ population. (D-H) Cytokine levels in CD8+/CD44+ T cells. Statistical significance was determined by unpaired *t-*test (\**p*<0.05, \*\**p*<0.01).

**Figure 7.**
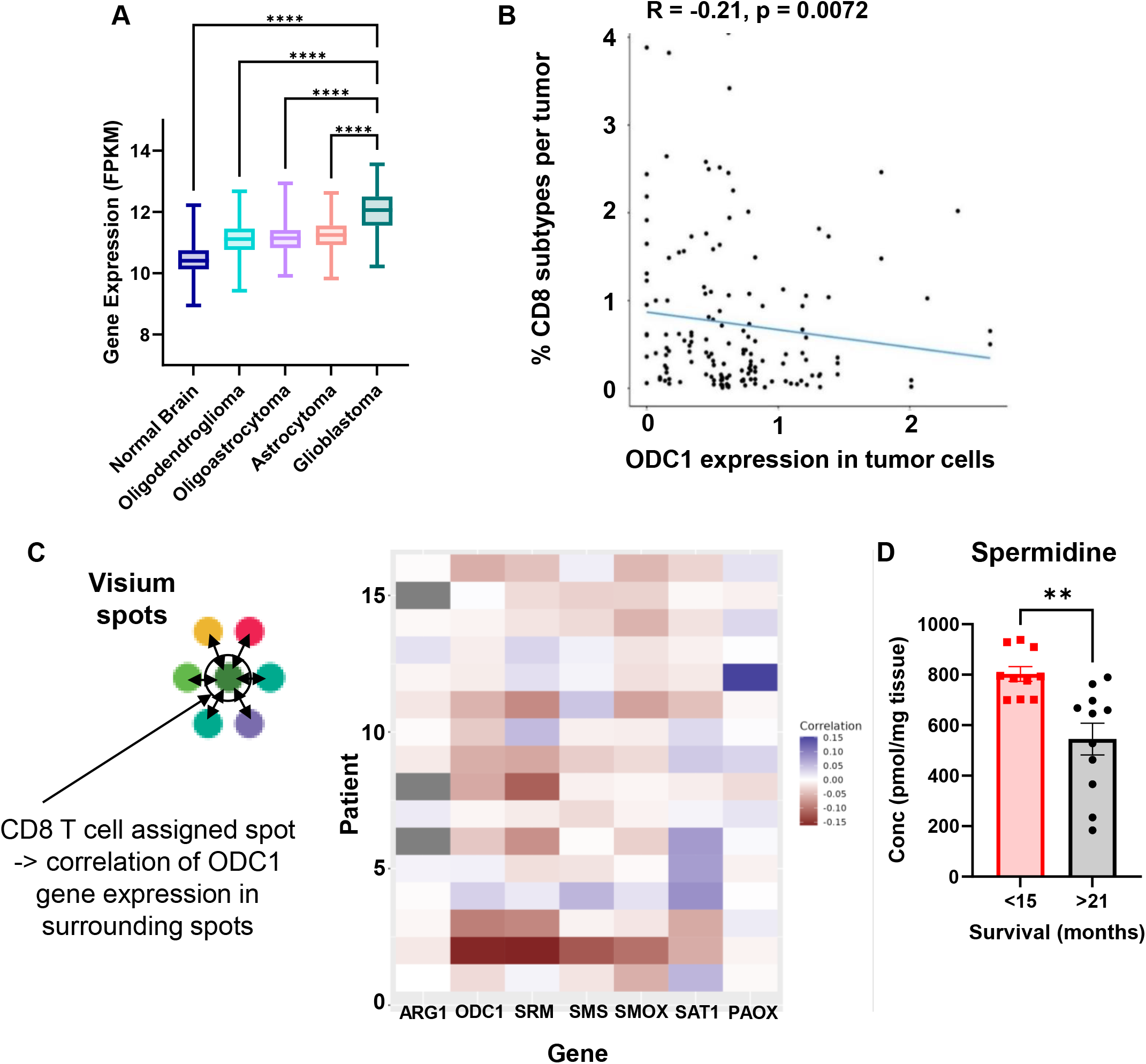
GBM patients have increased *ODC1* expression, and spermidine correlated with poorer prognosis. A) mRNA expression of *ODC1* from GTEX non-neoplastic and TCGA lower-grade gliomas and GBM tumor tissue. (B) Single-cell RNAseq correlation of *ODC1* expression in tumor cells and number of CD8+ cells in the tumor microenvironment. (C) Schematic of Visium single-cell analysis; heatmap showing that CD8+ T cells presence correlates with surrounding polyamine pathway gene expression by tumor cells. (D) Tumor tissue from primary resection of age-matched patients with longer survival compared to shorter survival; metabolites measured via LC-MS/MS. Statistical significance in (A) was determined by one-way ANOVA (****p<0.001). Statistical significance in (B) was determined by linear regression. Statistical significance in (D) was determined by unpaired *t-*test (\*\**p*<0.01). ARG1: arginase, ODC1: ornithine decarboxylase, SRM: spermidine synthase, SMOX: spermidine oxidase, SAT1: spermidine/ spermine acetyl transferase, PAOX: polyamine oxidase.

## Discussion

Here, we identify a new molecular mechanism through which GBM cells affect their surrounding microenvironment and drive a pro-tumorigenic state through direct depletion and impairment of T cells (**Fig. 8**). This immune alteration occurs via increased SPD in the tumor microenvironment and is driven by expression of ODC, the rate-limiting enzyme in the main polyamine biosynthesis pathway. These findings reinforce a model in which tumor cells secrete a host of factors to alter the immune microenvironment in their favor. Our findings show that SPD itself, either increased via exogenous addition or reduced via *ODC1* knockdown, did not alter intrinsic tumor growth but did impact cytotoxic T cells and Tregs. These results are similar to our previous observation in which GBM cancer stem cells secreted macrophage migration inhibitory factor, which supported myeloid-derived suppressor cell function but was dispensable for tumor cell growth^35^. It is worth noting that other studies have demonstrated an essential role for SPD in tumor cell growth, including in pediatric glioma and neuroblastoma. With respect to the differences between GBM and pediatric glioma in terms of SPD dependency, this could be due to inherent mutational landscapes and/or differential metabolic dependencies. Another possibility could be differing SPD levels between pediatric glioma and GBM cells, as previous observations in pediatric glioma were not directly compared to GBM models. It could be the case that GBM cells have an increased level of SPD at baseline compared to pediatric glioma cells; in this case, increasing spermidine would not elicit a pro-growth phenotype, and knockdown, which we employed here instead of complete knockout, would maintain a sufficient amount of SPD present to perpetuate cell growth.

**Figure 8.**
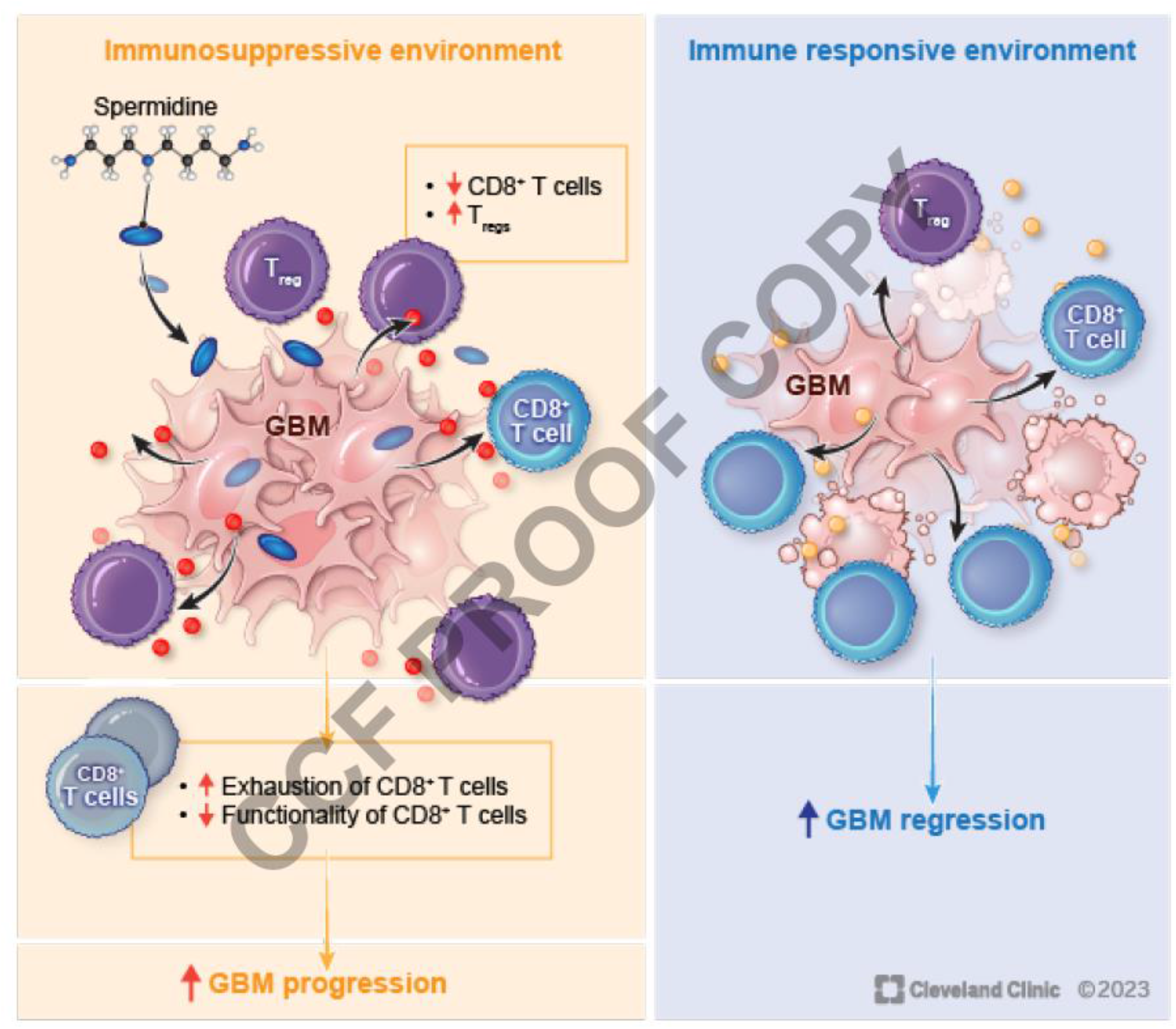
Summary Figure.

Our data support a model in which CD8+ T cells in the GBM microenvironment are more sensitive to changes in SPD compared to other immune cells. These findings are complementary to recent work in tumor-associated myeloid cells and may help explain why spermidine generates a pro-tumorigenic environment^22^, as it can increase immune suppression through enhancement of myeloid cells while concomitantly decreasing immune activation though the depletion and functional inhibition of cytotoxic T cells. Future studies would benefit from the direct comparison between these two pro-tumorigenic mechanisms to determine which population is more responsive to SPD, either directly or through other immune alterations.

While our studies focused on SPD, the polyamine family also contains the additional metabolites putrescine and spermine, as well as cadaverine, which is produced solely by bacteria. We observed that exogenous spermine administration does, to an extent, replicate the effects of SPD administration, resulting in a shortening of survival (*data not shown*). However, we did not observe a similar effect with putrescine administration (*data not shown*). There could be several reasons for the specificity of SPD compared to other polyamine family members. Although some polyamine functions are shared by all members, certain functions are driven mostly by a particular polyamine compared to the others. Cell necrosis and apoptosis are mediated by putrescine and SPD^20^. Another function that is more specific to SPD is inflammation reduction^36,37^. This correlates with the immune suppression we see in our studies as well as the characterization of GBM as a “cold tumor”^38^. While our studies focused on GBM, polyamines have been reported to have a pro-tumorigenic role in established tumors in other cancers – such as prostate and colorectal – and a tumor suppressive role at the initial stages in other tumors – namely melanoma and some types of breast cancer^39–43^. Therefore, our findings may be of interest to other tumor types.

Our studies leverage pre-clinical models to demonstrate that SPD can drive tumor growth in an immune-dependent manner and are consistent with other pediatric and adult brain tumor pre-clinical findings. Conceptually, these findings support the use of polyamine inhibitors for malignant brain tumors. However, current attempts to target these pathways via difluoromethylornithine, which is decarboxylated by ODC and binds to the enzyme, thus irreversibly inactivating it, have shown modest clinical efficacy^44^. This could partially be due to the ubiquitous nature of polyamines in the human body. Although this inhibitor blocks de novo biosynthesis of polyamines, uptake of polyamines secreted by other cells in the environment could help maintain tumor cell growth and sustain pressure on the immune response.

We should note that there are also limitations to our current study. The majority of our assessments are based in mouse models, and while we have some indication that SPD may function in humans in a manner similar to that of our pre-clinical models, additional interrogation of SPD and other polyamines in human tissue, CSF, and blood across a large cohort over tumor progression would be useful to determine the extent to which elevated SPD levels indicate immune suppression and poor prognosis. While our studies focused on lymphocyte changes, there are reports of a contribution by myeloid cells^22,45,46^, and together, these immune cell types could synergistically create a more pro-tumorigenic microenvironment. Focused studies interrogating both myeloid and lymphoid components will help clarify the effect of SPD on each immune lineage. As there is not one clear mechanism that accounts for the majority of cytotoxic T cell depletion and loss of functionality in a SPD-dependent manner, additional clarification is required to facilitate targeting strategies. Of note, while polyamines have been shown to impact T cell lineage specification via hypusination^47^, we did not observe an increase in hypusination in bone marrow-derived cells treated with SPD (*data not shown*), which could be due to many factors, including alternative pathway utilization. Finally, as our studies focused on polyamines produced by GBM cells in the tumor microenvironment, it should be noted that peripheral polyamines, including those originating from the gut microbiome, could also play a role in the overall immune response to GBM.

Our observations support a role for SPD in tumor microenvironment driving tumor growth, but there are also several unanswered questions based on these initial findings. We know that tumor cells produce higher levels of polyamines, but polyamines are also produced by other cells in the body and commensal gut microbes and are also found in the diet/taken in as part of the diet. Gut dysbiosis has been noted in many cancers, including GBM^48,49^, and there is a possibility that microbial reorganization in the gut could become skewed toward polyamine-producing strains, which would result in an increase in polyamines in the circulation and tumor microenvironment, thereby inducing immune suppression. Additionally, standard of care for GBM (surgical resection, radiation, chemotherapy) could affect both cellular production of polyamines as well as the gut microbiome, and this could result in altering the pool of polyamines or polyamine precursors available to cells in the tumor microenvironment. We show a reduction of cytotoxic functionality partially due to a reduction in CD8+ T cells and an increase in Tregs. The majority of immunotherapies rely on the presence of CD8+ T cells in the tumor microenvironment in order to augment their exhaustion and activation profiles^50^. Potentially, the inhibition of polyamine synthesis combined with the introduction of immunotherapies such as checkpoint inhibitors could increase the efficacy of immunotherapy in GBM. Finally, sex differences in the immune response have been noted in GBM, not only in localization of immune cells but also in their function and response to immunotherapies^26,51^, and the extent to which SPD and polyamines function in the context of sex differences is unclear. In our pre-clinical studies, we assessed males and females and observed no substantial sex differences; however, future therapeutic studies should consider sex as a biological variable given the above-mentioned reports. Taken together, our data highlight the communication between tumor cells and immune cells, which results in a favorable immune microenvironment for GBM growth and provides a function for SPD in the tumor microenvironment in facilitating this process.

## Materials and Methods

### Cell lines

The syngeneic mouse GBM cell line SB28 was a kind gift from Dr. Hideho Okada at University of California San Francisco, and GL261 cells were obtained from the Developmental Therapeutic Program at the National Cancer Institute. The CT-2A cell line was a kind gift from Prof. Misty Jenkins at the WEHI Australia. PC-3 human prostate cancer cells were obtained from Cleveland Clinic Lerner Research Institute. The patient-derived GBM model DI318 was derived at the Cleveland Clinic Lerner Research Institute, L1 was obtained from the University of Florida, and 3832 was obtained from Duke University. Human astrocytes were purchased from ScienCell. All cell lines were treated with 1:100 MycoRemoval Agent (MP Biomedicals) upon thawing and routinely tested for Mycoplasma spp. (Lonza). Mouse GBM cell lines and human prostate cancer cells were maintained in complete RPMI1640 (Media Preparation Core, Cleveland Clinic) supplemented with 10% FBS (Thermo Fisher Scientific) and 1% penicillin/streptomycin (Media Preparation Core, Cleveland Clinic). Human GBM lines, human astrocytes, and primary mouse microglia and astrocytes were maintained in complete DMEM:F12 (Media Preparation Core, Cleveland Clinic) supplemented with 1% penicillin/streptomycin, 1X N-2 Supplement (Gibco), and EGF/FGF-2. All cells were maintained in humidified incubators held at 37°C and 5% CO2 and not grown for more than 20 passages.

### Mice

All animal procedures were performed in accordance with the guidelines and protocols approved by Institutional Animal Care and Use Committee (IACUC) at the Cleveland Clinic and by the Walter and Eliza Hall Institute Animal Ethics Committee. C57BL/6 (RRID: IMSR_JAX:000664) and RAG1^-/-^ (B6.129S7-Rag1tm1Mom/J; RRID: IMSR_JAX:002216) mice were purchased from the Jackson Laboratory as required. NSG (NOD.Cg-Prkdc^scid^Il2rg^tm1Wjl^/SzJ) mice were obtained from the Biological Research Unit (BRU) at Lerner Research Institute, Cleveland Clinic. All animals were housed in a specific-pathogen-free facility of the Cleveland Clinic BRU with a light-dark period of 12 h each. All animals were maintained on a control diet to minimize/normalize polyamines consumed via the diet (Research Diets, D12450J).

For tumor implantation, 5–8-week-old mice were anesthetized, fit to a stereotactic apparatus, and intracranially injected with 10,000-25,000 tumor cells in 5 µl RPMI-null media into the left hemisphere approximately 0.5 mm rostral and 1.8 mm lateral to the bregma with 3.5 mm depth from the scalp. In CT-2A experiments, 10,000 tumor cells were injected 1 mm lateral, 1 mm anterior with 2.5 mm depth. In some experiments, 5 µl null media was injected into age- and sex-matched animals for sham controls. Animals were monitored over time for the presentation of neurological and behavioral symptoms associated with the presence of a brain tumor. Biological sex is indicated for each study.

In some experiments, mice were treated with 50 mg/kg SPD (Sigma, cat# S0266) diluted in 0.9% saline or 0.9% saline control intraperitoneally starting from 7 days post-tumor implantation; mice received 3 injections per week until endpoint.

## Isolation of *ex vivo* mouse cells for *in vitro* testing

### Microglia and Astrocytes

Primary mouse microglia and astrocytes were isolated and cultured from D0-D1 wild-type B6 pup brains, as previously described^52^.

***CD8^+^/CD4+ T cells*** were isolated from splenocytes of 8–12-week-old mice using magnetic bead isolation kits (Stemcell Technology). Isolated CD8^+^ T cells were cultured in the presence of recombinant human IL-2 (100 U/ml, PeproTech) and anti-CD3/CD28 Dynabeads (Thermo Fisher Scientific) for 3-4 days before flow cytometry studies. ***T regulatory cells*** were cultured from CD4+ T cells and induced with IL-2 (100 U/ml, PeproTech), anti-CD3/CD28 Dynabeads (Thermo Fisher Scientific), and TGFβ (5 ng/ml, PeproTech). For proliferation studies, T cells were stained with 1:1000 CellTrace Violet (Invitrogen) prior to culturing

### MDSCs

Bone marrow was isolated from the femur and tibia of 8- to 12-week-old mice. Two million bone marrow cells were cultured in 6-well plates in 2 mL RPMI/10% FBS supplemented with 40 ng/mL GM-CSF and 80 ng/mL IL13 (PeproTech) for 3 to 4 days. Cells were stained for viability, blocked with Fc receptor inhibitor and stained with a combination of CD11b, Ly6C, and Ly6G for sorting of MDSC subsets (mMDSCs: CD11b^+^Ly6C^+^Ly6G^−^ vs. gMDSCs: CD11b^+^Ly6C^−^Ly6G^+^) and the control population (CD11b^+^Ly6C^−^Ly6G^−^) using a BD FACSAria II (BD Biosciences).

### Cell viability and functionality assays

The cell models described above were treated with varying concentrations of SPD in DMSO/PBS or equivalent vehicle in respective complete media. At the time points described in the corresponding figure legends, single-cell suspensions were combined with an equal volume of 0.4% Trypan Blue (Thermo) and counted using a TC20 Automatic Cell Counter (Bio-Rad). Alternatively, an equal volume of CellTiter-Glo Luminescent Cell Viability Assay (Promega) was added to treated cells, and viability was measured via luminescence on a VICTOR Nivo multimode plate reader (PerkinElmer).

To measure cell death and apoptosis of CD8+ T cells treated *in vitro* with SPD, FITC-labeled annexin V (BioLegend) and DRAQ7 (Invitrogen) were added in accordance with the manufacturer’s protocols. To measure intracellular pH levels, CD8+ T cells were labeled with pHrodo Red (ThermoFisher) according to the manufacturer’s protocol. Samples were run on an LSR Fortessa flow cytometer (BD Biosciences) with a minimum of 10,000 events collected. Single cells were gated, and the percentages of annexin V- and/or DRAQ7-positive cells were determined. For pHrodo Red-labeled cells, high and low gates were used to determine intracellular acidic and neutral pH based on gMFI (geometric mean fluorescence intensity – a measure of the shift in fluorescence intensity of a population of cells). For intracellular cytokine detection, cells were stimulated using Cell Stimulation Cocktail plus protein transport inhibitor (eBioscience) in complete RPMI for 4 hours. After stimulation, cells were subjected to the flow cytometry staining procedures described below.

### Liquid chromatography-mass spectrometry quantification of polyamine metabolites Sample preparation

Plasma and tissue samples for polyamine quantitation were processed as previously described for serum samples, with minor modifications as below^53^.

Twenty microliters of plasma was aliquoted into a 12 x 75 mm glass tube and mixed with 5 μl internal standard mix consisting of [2H5]ornithine, [13C6]arginine, [2H8]spermine, [2H8]spermidine, [13C4]putrescine and [2H3]acisoga in water with a concentration (in μM) of 400, 400, 10, 10, 10 and 0.5, respectively. Then 5 μl of 1 M sodium carbonate (pH 9.0) and 10 μl isobutyl chloroformate were added to derivatize polyamines. Then 0.5 ml diethyl ether was added to extract the derivatized product. All the stable isotope internal standards were purchased from Cambridge Isotope Lab or CDN Isotopes.

For the tissue samples, approximately 20 mg brain tissue was mixed with 5 μl of the above internal standard mix in a 2 ml Eppendorf tube with 400 μl H2O, followed by homogenization in a tissue homogenizer (Qiagen) with a metal bead (Qiagen #69997) added. The homogenate was spun down at 20,000 x g at 4°C for 10 minutes. Supernatant (200 µl) was transferred to a clean 12 x 75 mm glass tube, and 50 μl of 1 M sodium carbonate (pH 9.0) and 100 μl isobutyl chloroformate were added to derivatize polyamines. Then 2 ml diethyl ether was added to extract the derivatized product. The diethyl ether extract was dried under N2 and resuspended in 50 μl of 1:1 0.2% acetic acid in water:0.2% acetic acid in acetonitrile and transferred to a mass spectrometer with plastic insert for LC/MS assay.

### Liquid chromatography–mass spectrometry (LC/MS) assay

Supernatants (5 μl) were analyzed by injection onto a Cadenza CD-C18 Column (50 x 2 mm, Imtaket) at a flow rate of 0.4 ml/min using a Vanquish LC autosampler interfaced with a Thermo Quantiva mass spectrometer. A discontinuous gradient was generated to resolve the analytes by mixing solvent A (0.2% acetic acid in water) with solvent B (0.2% acetic acid in acetonitrile) at different ratios starting from 0% B to 100% B. The mass parameters were optimized by injection of individual derivatized standard or isotope labeled internal standard individually. Nitrogen (99.95% purity) was used as the source, and argon was used as collision gas. Various concentrations of nonisotopically labeled polyamine standard mixed with internal standard mix undergoing the same sample procedure was used to prepare calibration curves.

### Immunophenotyping by flow cytometry

At the indicated time points, a single-cell suspension was prepared from the tumor-bearing left hemisphere by enzymatic digestion using collagenase IV (Sigma) and DNase I (Sigma). Digested tissue was filtered through a 70-µm cell strainer, and lymphocytes were enriched by gradient centrifugation using 30% Percoll solution (Sigma). Cells were then filtered again with a 40-µm filter. Cells were stained with LIVE/DEAD Fixable stains (Thermo Fisher) on ice for 15 min. After washing with PBS, cells were resuspended in Fc receptor blocker (Miltenyi Biotec) diluted in PBS/2% BSA and incubated on ice for 10 min. For surface staining, fluorochrome-conjugated antibodies were diluted in Brilliant Buffer (BD) at 1:100 – 1:250, and cells were incubated on ice for 30 min. After washing with PBS-2% BSA buffer, cells were then fixed with FOXP3/Transcription Factor Fixation Buffer (eBioscience) overnight. For intracellular staining, antibodies were diluted in FOXP3/Transcription Factor permeabilization buffer (perm buffer) at 1:250-1:500, and cells were incubated at room temperature for 45 min. For intracellular cytokine detection, cells were stimulated using Cell Stimulation Cocktail plus protein transport inhibitor (eBioscience) in complete RPMI for 4 hours. After stimulation, cells were subjected to the staining procedures described above. Stained cells were acquired with a BD LSR Fortessa (BD) or Aurora (Cytek) and analyzed using FlowJo software (v10, BD Biosciences).

### Reagents

For immunophenotyping in mouse models, the following fluorophore-conjugated antibodies at concentrations of 1:250-1:500 were used: CD11b (M1/70, Cat# 563553), CD11c (HL3, Cat# 612796), CD3 (145-2C11, Cat# 564379), and CD44 (IM7, Cat# 612799) from BD biosciences. CTLA4 (UC10-4B9, Cat# 106312), PD1 (29F.1A12, Cat# 135241), B220 (RA3-6B2, Cat# 103237), Ki-67 (11Fb, Cat# 151215), TIM3 (RMT3-23, Cat# 119727), I-A/I-E (M5/114.15.2, Cat# 107606), CD45 (30-F11, Cat# 103132), LAG3 (C9B7W, Cat# 125224), NK1.1 (PK136, Cat# 108716), CD4 (GK1.5, Cat# 100422), CD8 (6206.7, Cat# 100712), granzyme B (QA18A28, Cat# 396413), TNFα (MP6-XT22, Cat# 506329), and IFNγ (XMG1.2, Cat# 505846) were obtained from BioLegend. Anti-Foxp3 (FJK-16s, Cat# 12-5773) antibody was obtained from eBioscience.

### Stable transduction with lentiviral shRNA

Lentifect Ultra-Purified Lentiviral Particles targeting mouse *ODC1* and an associated non-targeted control lentiviral particle were purchased from Genecopoiea. Prior to transfection, mouse glioma cells were grown to ∼70% confluency on tissue-culture treated plates. Lentivirus was added to and incubated with the cells for 24 h, followed by a change to fresh media. Selection was then initiated with puromycin (ThermoFisher). Transfected cells were incubated in media with 3 µg/ml puromycin for 48 h. Stably transfected cells were maintained in their regular media plus puromycin at 1 µg/ml. Knockdown was verified via RT-qPCR.

### Real-time quantitative PCR

Total RNA was isolated using an RNeasy mini kit (Qiagen), and cDNA was synthesized with qSCRIPT cDNA Super-mix (Quanta Biosciences). qPCR reactions were performed using Fast SYBR-Green Mastermix (Thermo Fisher Scientific) on an Applied Biosystems QuantStudio 3 Real-Time PCR system. The threshold cycle (Ct) value for each gene was normalized to the expression levels of *Gapdh*, and relative expression was calculated by normalizing to the delta Ct value of mouse astrocytes, unless otherwise described. Primer sequences were obtained from PrimerBank or previously published papers and are listed in Table S1 (mouse).

### TCGA and GTEX data analysis

Clinical and mRNA expression data for the IDH-wildtype subset of GBM cohort and lower-grade glioma cohorts of TCGA were downloaded from the GlioVis portal (http://gliovis.bioinfo.cnio.es); GBM and normal brain cohorts of GTEX were downloaded from the GTEX portal (https://gtexportal.org/home/).

#### Analysis of single-cell RNAseq data from Ruiz-Moreno et al

Publicly available dataset GBmap was utilized and analyzed using Seurat v4.0 (https://www.biorxiv.org/content/10.1101/2022.08.27.505439v1.full.pdf, Ruiz-Moreno et al., biorxiv, 2022). The Core GBmap data was downloaded, which comprises 338,564 total cells harmonized from 16 different studies. Briefly, the authors used a semi-supervised neural network model to integrate the data and used any additional data to classify cell type. Furthermore, they used gene modules to further categorize cell types. The Seurat rds file was downloaded, and the cell type annotations determined by GBmap were used. The average *ODC1* expression per sample was calculated using Seurat’s AverageExpression function. CD8 cytotoxic, CD8 EM, and CD8 NK sig cells were aggregated to represent the CD8-expressing cells per tumor. For each sample, the percentage of CD8-expressing cells was calculated, using the total number of cells per sample as the denominator. A Spearman correlation was calculated and plotted in Figure 7.

#### Analysis of Visium spatial transcriptomics data from Ravi et al

Processed data were downloaded from https://doi.org/10.5061/dryad.h70rxwdmj. Deconvolution of spots as described in Ravi et al. were obtained from the authors upon request. We calculated the correlation between the gene expression of interest in each spot and the average proportion of estimated CD8+ T cells in all adjacent spots using a simple Pearson correlation.

### MALDI-TOF spatial analysis

Flash-frozen tissue was sectioned at a thickness of 10 µm directly onto Indium Tin Oxide (ITO)-coated glass slides. Frozen sections were dried in a freeze dryer (MODULYOD, Thermo Electron Corporation) for 30 min, followed by collection of optical images using the light microscope embedded in the MALDI-TOF MSI instrument (iMScope^TM^ QT) prior to matrix application. α-cyano-4-hydroxycinnamic acid (CHCA, P# C2020) was purchased from Sigma-Aldrich, Germany. Matrix deposition was performed by two-step deposition method using iMLayer for sublimation and iMLayer AERO (Shimadzu, Japan) for matrix spraying. The thickness of the vapor-deposited matrix was 0.7 µm, and the deposition temperature was 250°C. For CHCA matrix spraying, 8 layers of 10 mg/mL CHCA in acetonitrile/water (50:50, v/v) with 0.1% trifluoroacetic acid solution were used. The stage was kept at 70 mm/sec with 1 sec dry time at a 5 cm nozzle distance and pumping pressure kept constant at 0.1 and 0.2 MPa, respectively. MALDI-TOF experiments were performed using an iMScopeTM QT instrument (Shimadzu, Japan). The instrument is equipped with a Laser-diode-excited Nd:YAG laser and an atmospheric pressure MALDI. Data were collected at 10 µm spatial resolution with positive polarity.

### Bulk RNA sequencing

Normal and tumor regions were dissected from flash-frozen tissue, ground in liquid nitrogen and RNA extracted using the RNeasy RNA extraction kit (Qiagen 74104). TruSeq libraries (TruSeq RNA Library Prep v2, Illumina) were sequenced on the NextSeq System (Illumina) to produce 132 bp single-end reads.

### GBM patient samples

Frozen GBM specimens were collected by the Cleveland Clinic Rose Ella Burkhardt Brain Tumor and Neuro-Oncology Center after obtaining written informed consent from the patients. The studies were conducted in accordance with recognized ethical guidelines and approved by the Cleveland Clinic Institutional Review Board (IRB 2559).

### Statistical analysis

GraphPad Prism (Version 10, GraphPad Software Inc. RRID:SCR_002798) software was used for data presentation and statistical analysis. Unpaired or paired *t* test or one-way/two-way analysis of variance (ANOVA) was used with a multiple comparison test as indicated in the figure legends. Survival analysis was performed by log-rank test. *p*-value <0.05 was considered significant (\**p*<0.05, \*\**p*<0.01, \*\*\**p*<0.001).

## Data availability statement

All data generated in this study are available upon request from the corresponding author, Dr. Justin D. Lathia).

## Author’s contributions

Conception and design: K.E.K., J.L., D.B, J.D.L.

Development of methodology: K.E.K., J.L., D.B., Z.W.

Acquisition of data: K.E.K., J.L., D.B., J.B., S.D., E.W., S.Z.W., T.L., L.F., V.N.

Analysis and interpretation of data: K.E.K., J.L., D.B., E.S.H., J.V., T.L., L.F., V.N., S.F., S.B., J.W. J.D.L.

Writing, review: J.L., D.J.S., O.R., J.Y., S.H., J.M.B., D.B., J.D.L.

Administrative, technical, or material support: S.J., M.M., M.M.G., D.J.S., J.D.L.

Study supervision: J.D.L.

## Acknowledgements

We thank the members of the Lathia laboratory for insightful discussions. We are grateful to Drs. Jason Miska (Northwestern University) and Sameer Agnihotri (University of Pittsburgh) for their critical feedback. We greatly appreciate the editorial assistance of Dr. Erin Mulkearns-Hubert (Cleveland Clinic) and illustrative work of Ms. Amanda Mendelsohn from the Center for Medical Art and Photography at the Cleveland Clinic. We would like to acknowledge technical help from the Cleveland Clinic Flow Cytometry Core. This work is supported by National Institutes of Health grants F31 CA264849 (K.E.K.), T32 GM088088 (K.E.K), R35 NS127083 (J.D.L), P01 CA245705 (J.D.L.), and K99 CA248611 (D.B.). This work was also supported by the American Brain Tumor Association (J.D.L.), Case Comprehensive Cancer Center (J.D.L.), and the Cleveland Clinic/ Lerner Research Institute (J.D.L.). Additionally, this work was financially supported in part through the authors’ membership of the Brain Cancer Centre (T.L., L.F., S.F., S.A.B., J.R.W.), support from Carrie’s Beanies 4 Brain Cancer, a Perpetual Philanthropic Grant (IPAP20221259 to S.A.B., S.F. and J.R.W.), a Cancer Council Victoria Venture Grant (VG2022 to S.A.B., S.F. and J.R.W.), and through Victorian State Government Operational Infrastructure Support and Australian Government NHMRC Independent Research Institutes Infrastructure Support Scheme (IRIISS). Support from the Victorian Cancer Agency Mid-Career Research Fellowship (MCRF22003 to S.A.B.), and a National Health and Medical Research Council of Australia (NHMRC) Ideas Grant (GNT1184421 to S.F.).

## Disclosures

The authors declare no competing interests.

---

**Supplementary Figure 1.**
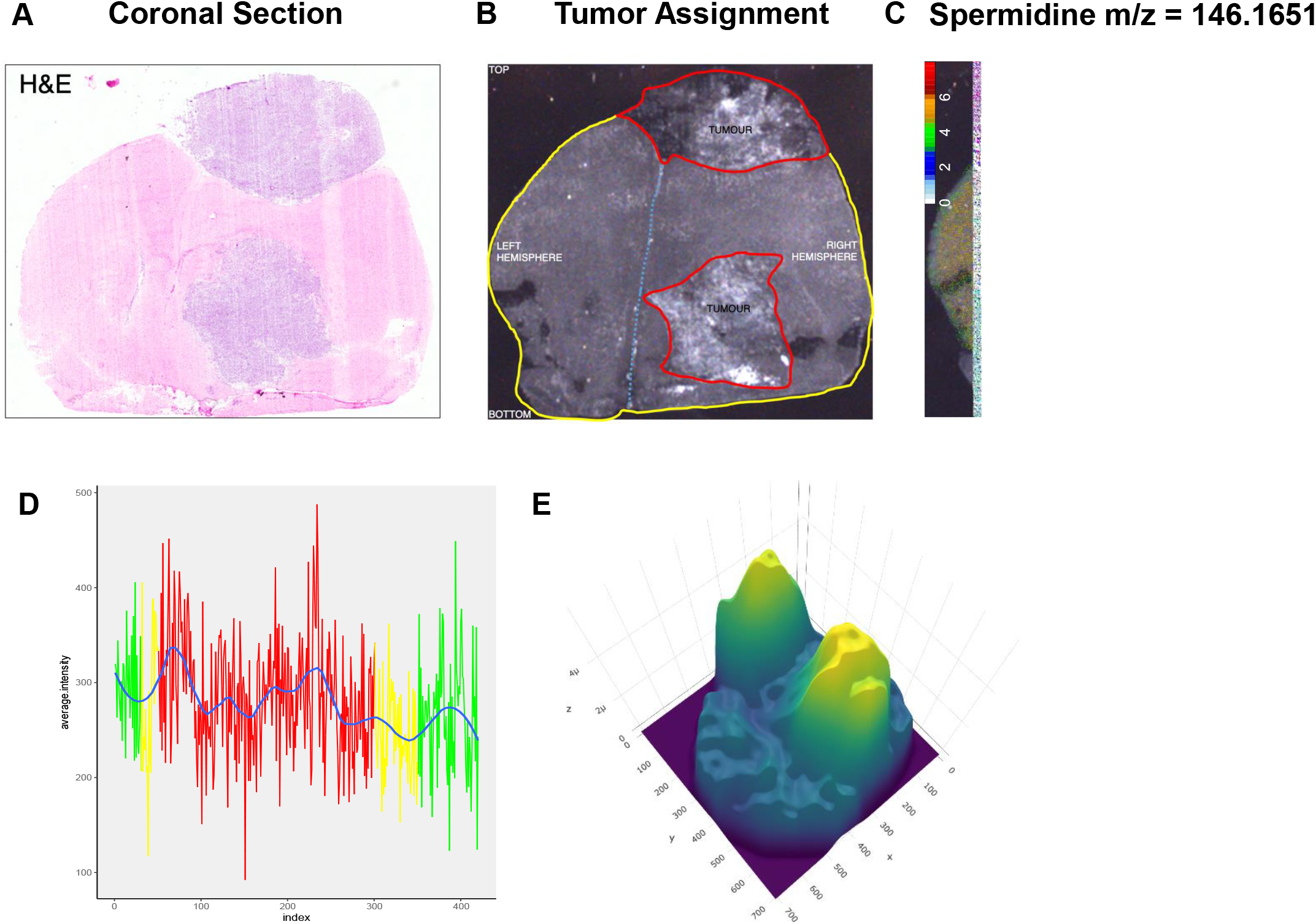
Spatial MALDI-TOF analysis of mouse tumors shows high SPD production in the tumor microenvironment. (A-C) After intracranial transplantation with syngeneic GBM cells (CT-2A), the mouse brain was removed and sectioned for staining and mass spectrometry. (A) H&E staining of brain tissue. (B) Identification and delineation of tumor areas in brain tissue. (C) SPD levels in the tumor and surrounding brain with corresponding intensity scale. (D) Line plot shows the average intensity of SPD across tumor (red rectangle), border (yellow rectangle) and adjacent normal tissue (green rectangle). (E) 3D plot of the estimated density of abundance of SPD.

**Supplementary Figure 2.**
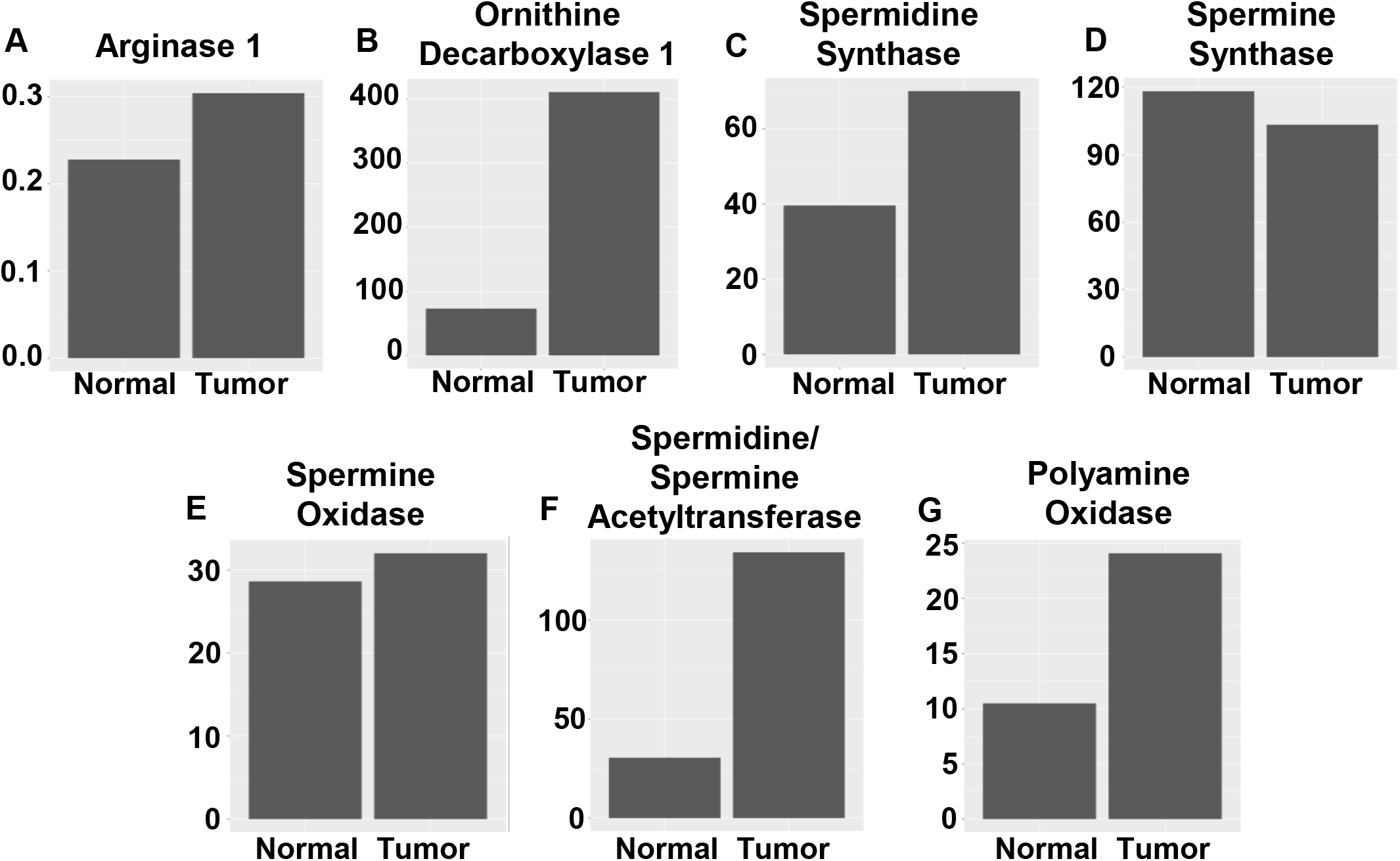
Bulk RNAseq data highlights increased polyamine pathway enzymes. (A-E) Tumor tissue and surrounding brain tissue from mice implanted with syngeneic GBM cells (CT2A) were processed via bulk RNA sequencing; expression of genes encoding enzymes of the main *de novo* polyamine synthesis pathway was analyzed.

**Supplementary Figure 3.**
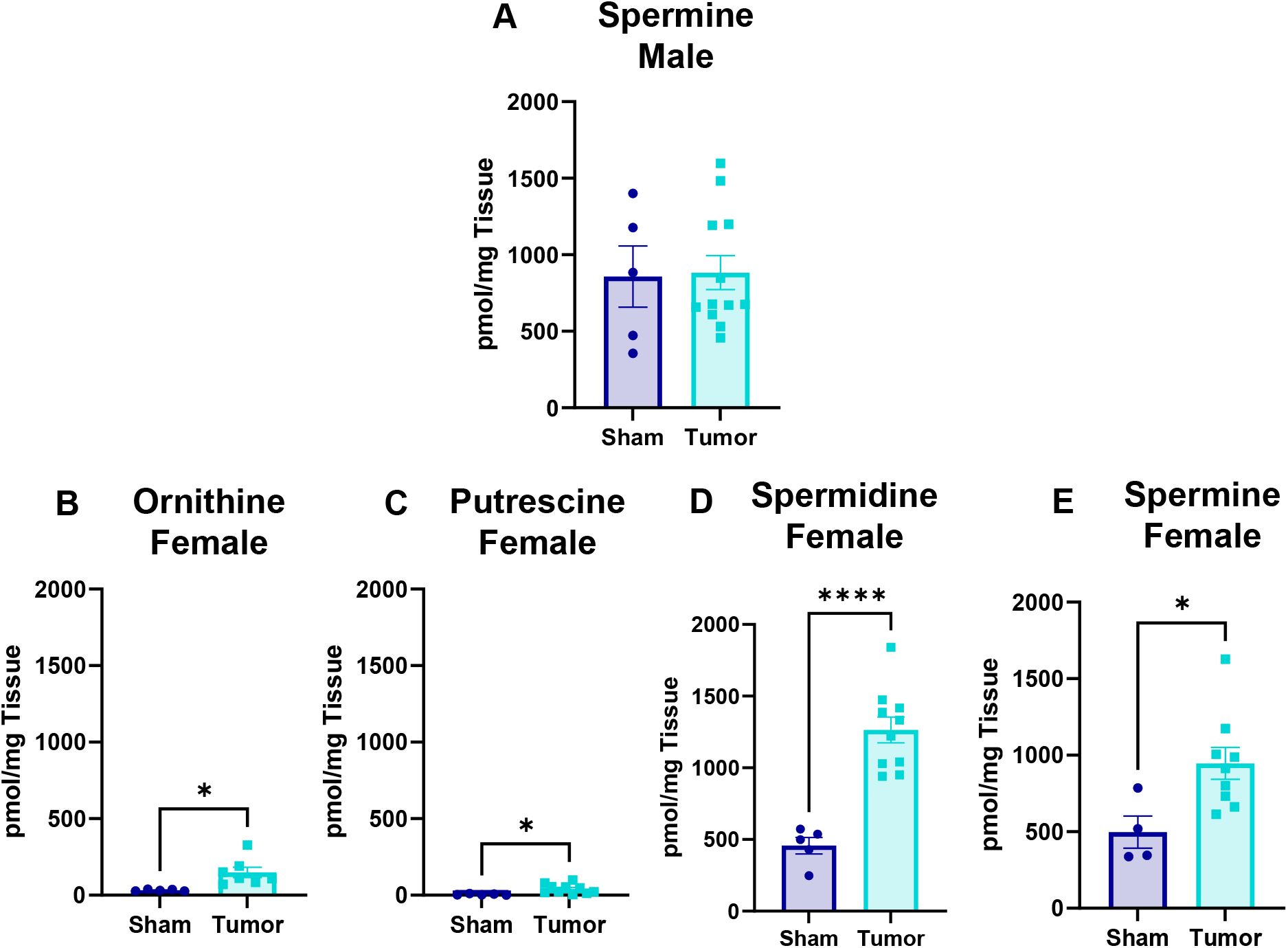
SPD levels are increased in females in a mouse GBM model. LC-MS/MS was performed on tumors removed from B6 mice 17 days after intracranial injection of mouse GBM cell line (25K/injection GL261). (A) Tumor levels of spermine in males. (B-E) Polyamine measurements in female mice. Statistical significance was determined by unpaired *t-*test (\**p*<0.05, \*\*\*\**p*<0.0001).

**Supplementary Figure 4.**
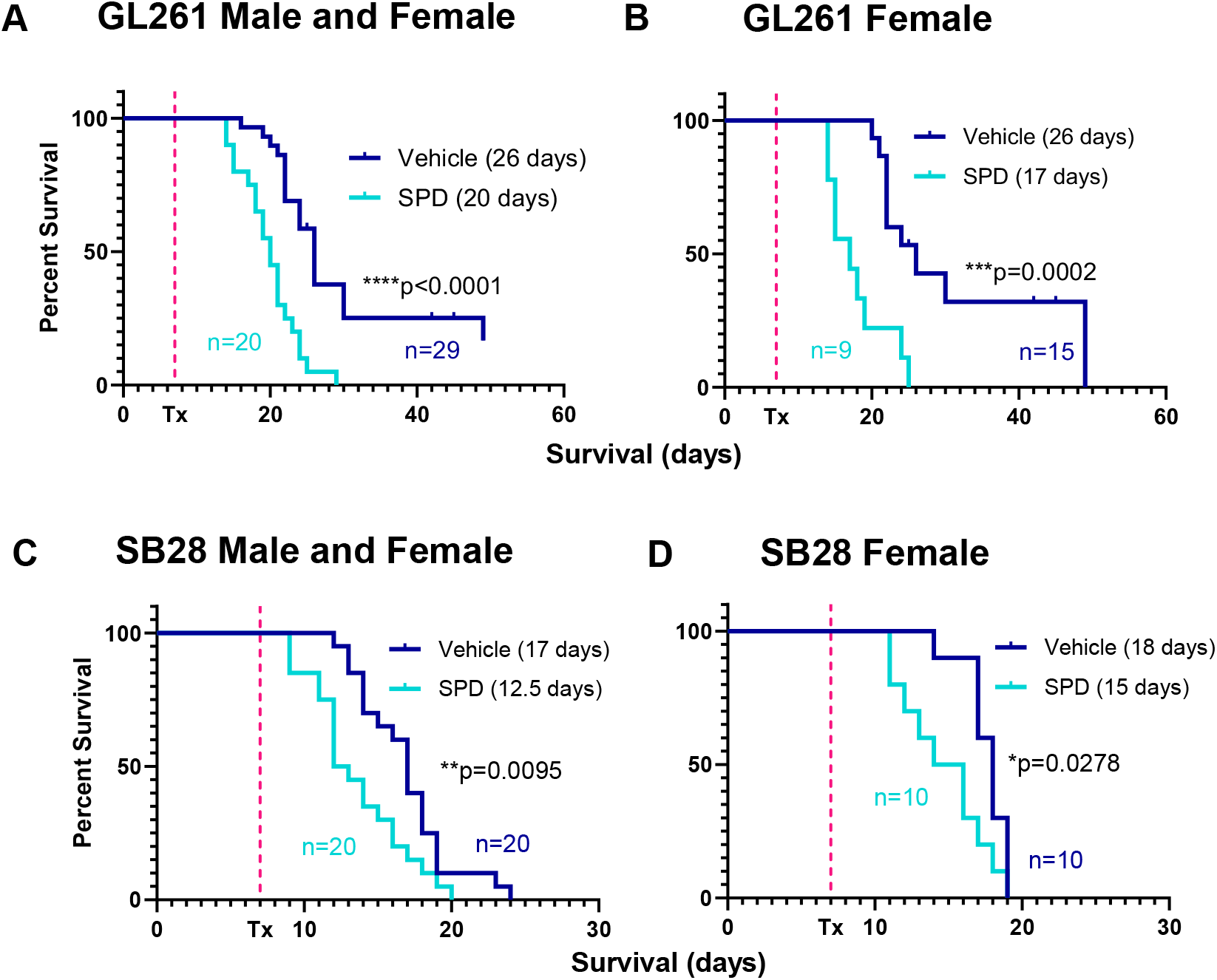
Exogenous SPD treatment of both male and female mice reduces survival. (A-D) Survival analysis was performed after intracranial injection of mouse GBM cell lines (25K/injection GL261, 20K/injection SB28) in B6 mice. Median survival days and number of animals are indicated in the figure. Data combined from three independent experiments. Statistical significance was determined by log-rank test, considering *p*-value <0.05 to be significant.

**Supplementary Figure 5.**
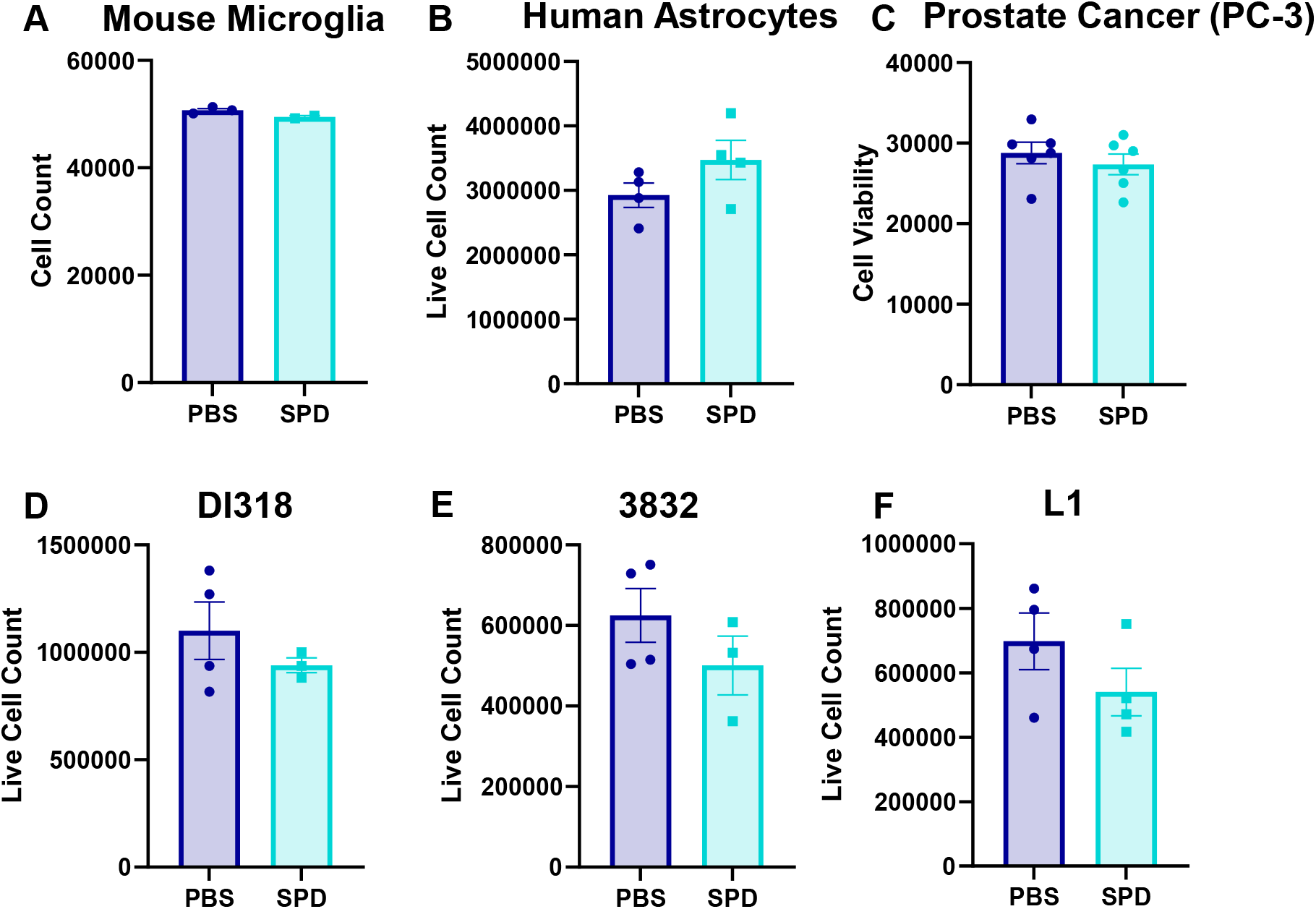
SPD does not affect viability of resident brain cells or human tumor cells *in vitro*. (A-F) Cells treated with physiological levels of SPD or vehicle *in vitro*. (A-B) Resident brain cells. (C) Human prostate cancer cells. (D-F) Human-derived GBM models.

**Supplementary Figure 6.**
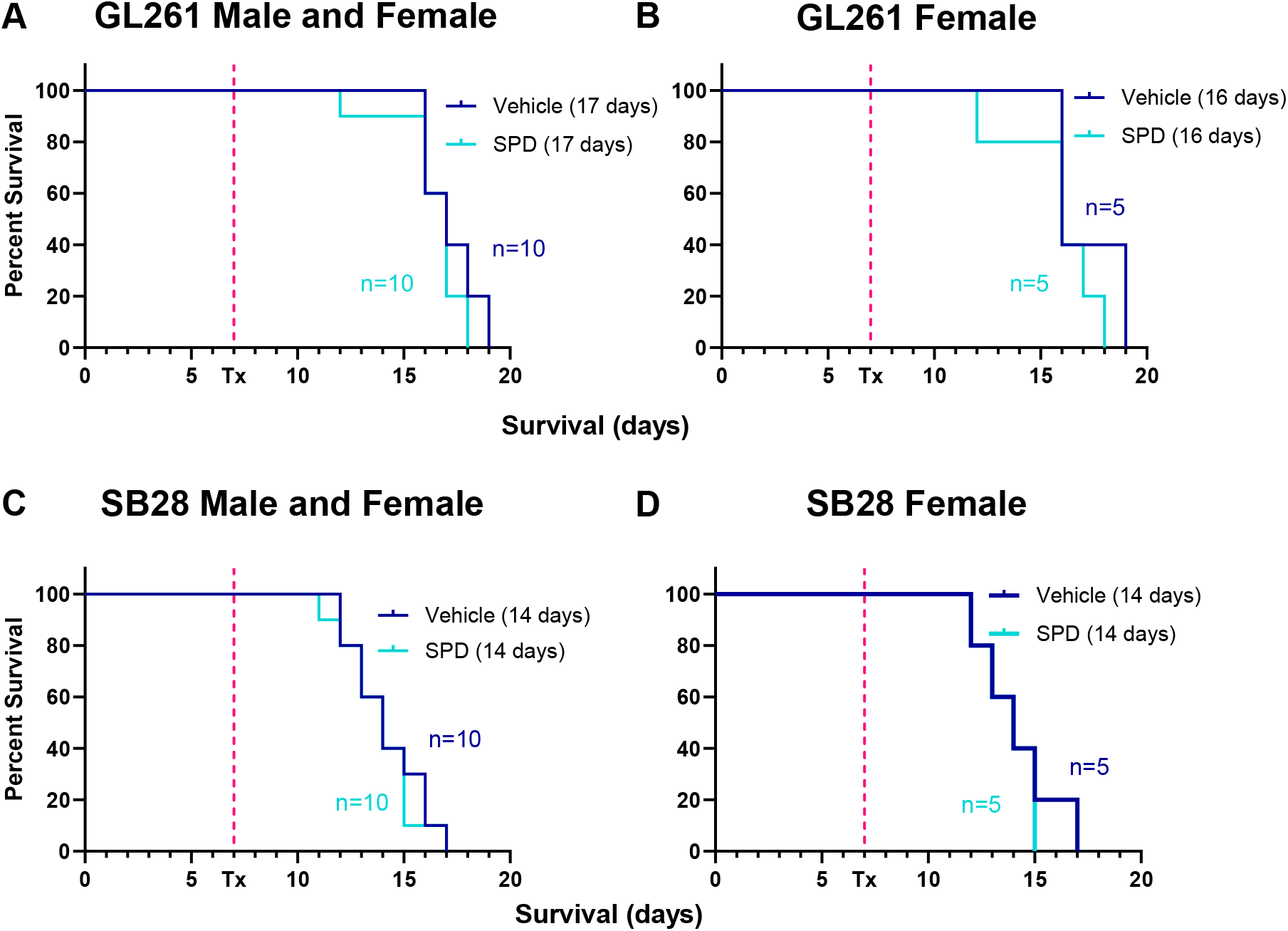
SPD interacts with the immune system to drive GBM progression in both male and female mice. (A-D) Survival analysis was performed after intracranial injection of mouse GBM cell lines (25K/injection GL261, 20K/injection SB28) in immunocompromised NSG mice. Median survival days and number of animals are indicated in the figure. Statistical significance was determined by log-rank test, considering *p*-value <0.05 to be significant.

**Supplementary Figure 7.**
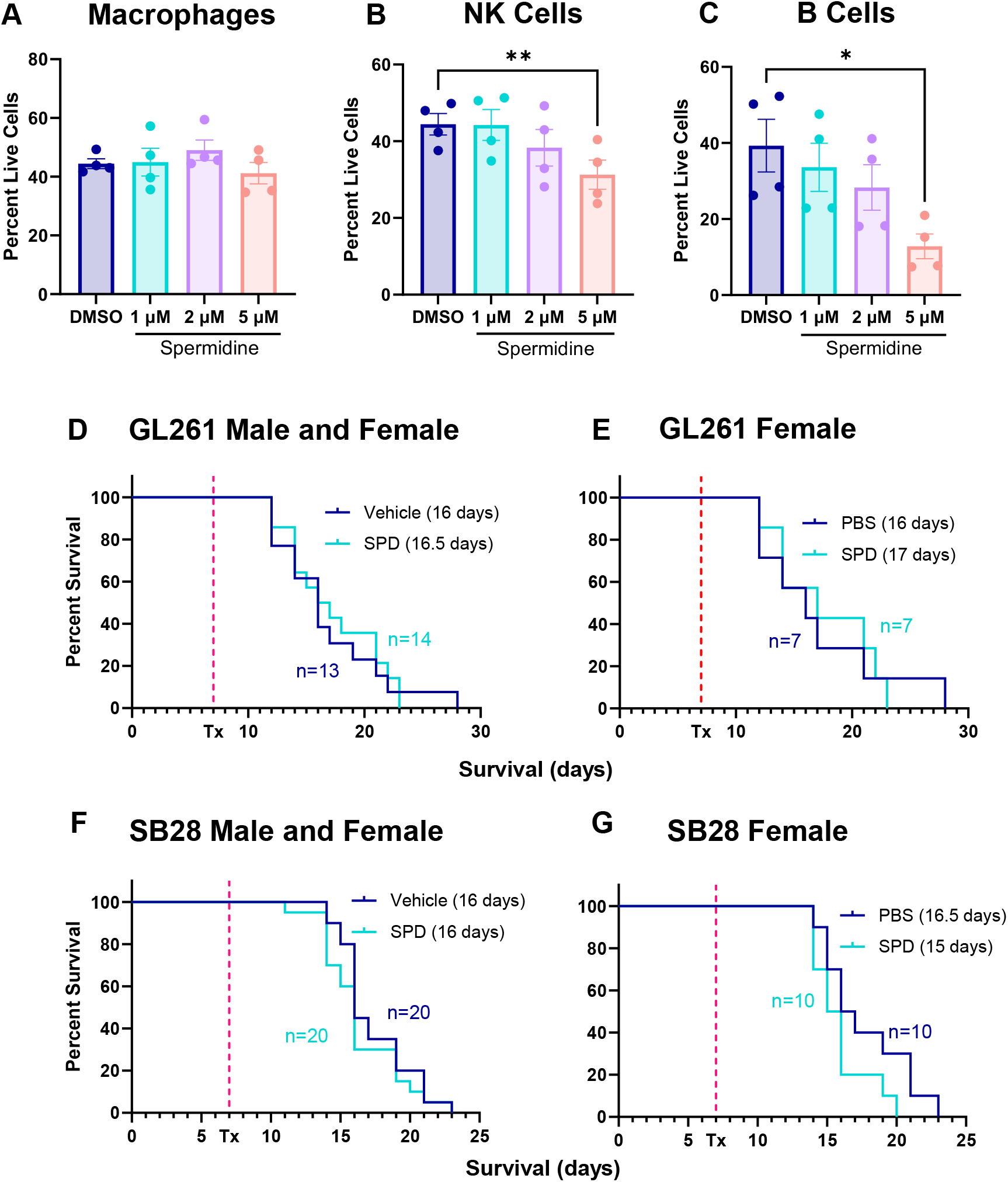
Lymphocyte subsets are affected by SPD. (A-C) Splenocyte-derived lymphocyte subsets were treated with physiological levels of SPD *in vitro*. (D-G) Survival analysis was performed after intracranial injection of mouse GBM cell lines (25K/injection GL261, 20K/injection SB28) followed by 50 mg/kg IP SPD treatment or PBS vehicle in Rag1 knockout mice. Median survival days and number of animals are indicated in the graph. Data combined from two independent experiments. Statistical significance for (A-C) was determined by one-way ANOVA (\**p*<0.05, \*\**p*<0.01). Statistical significance for (D-G) was determined by log-rank test, considering *p*-value <0.05 to be significant.

**Supplementary Figure 8.**
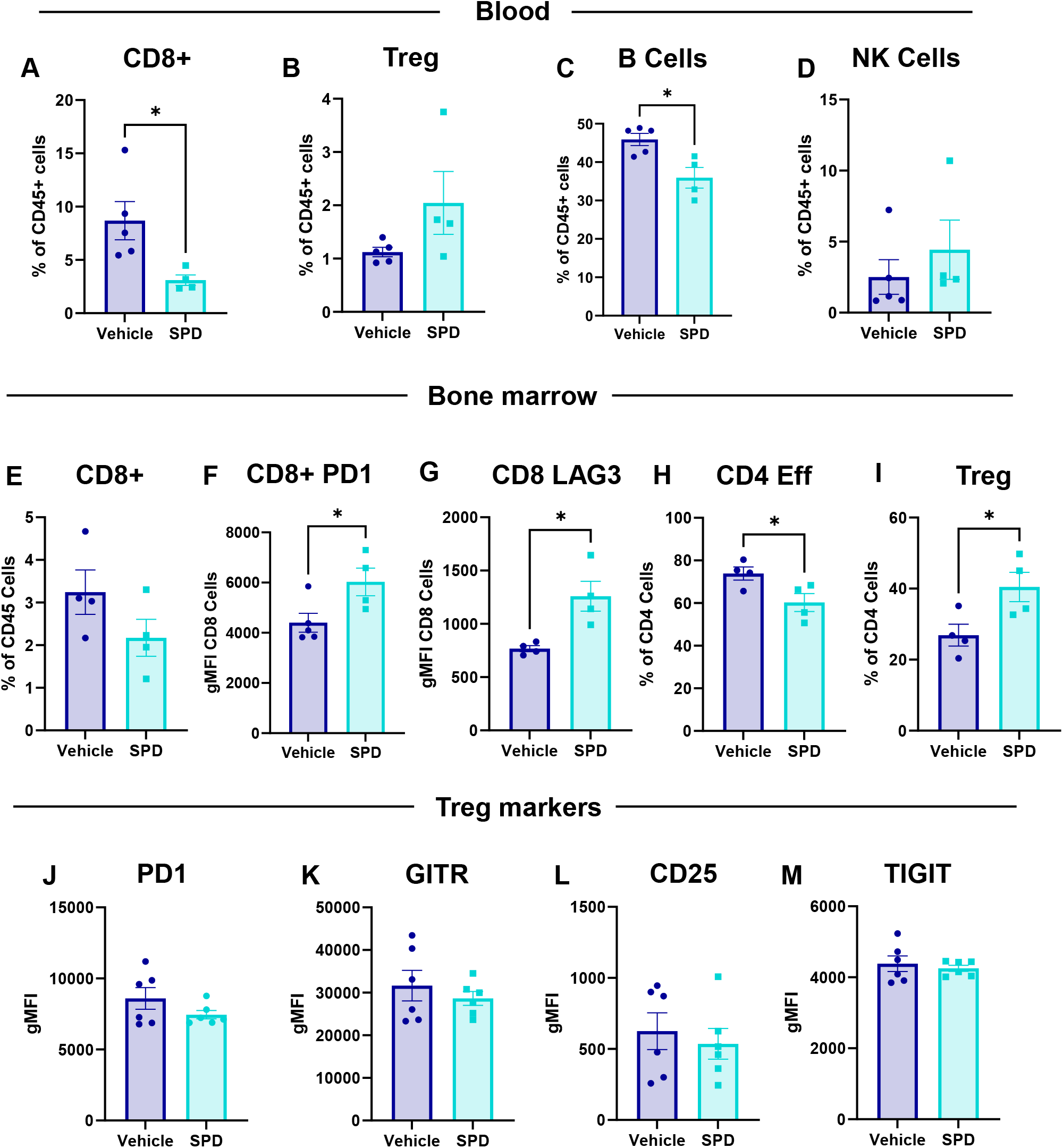
Effects of SPD on other immune compartments. After intracranial injection of mouse GBM cell line SB28 (20K/injection) into male B6 mice followed by 50 mg/kg SPD IP treatment or PBS vehicle, the tumor-bearing hemisphere was collected and processed for flow cytometry immune phenotyping. (A-D) Frequency of various immune cell types in blood. (E) Proportion of CD8+ T cells in bone marrow. (F-G) Exhaustion markers of CD8+ T cells. (H-I) Proportion of effector cells and Tregs in CD4+ population. (J-M) Treg markers. Statistical significance for (A-M) was determined by unpaired *t-*test (\**p*<0.05).

**Supplementary Figure 9.**
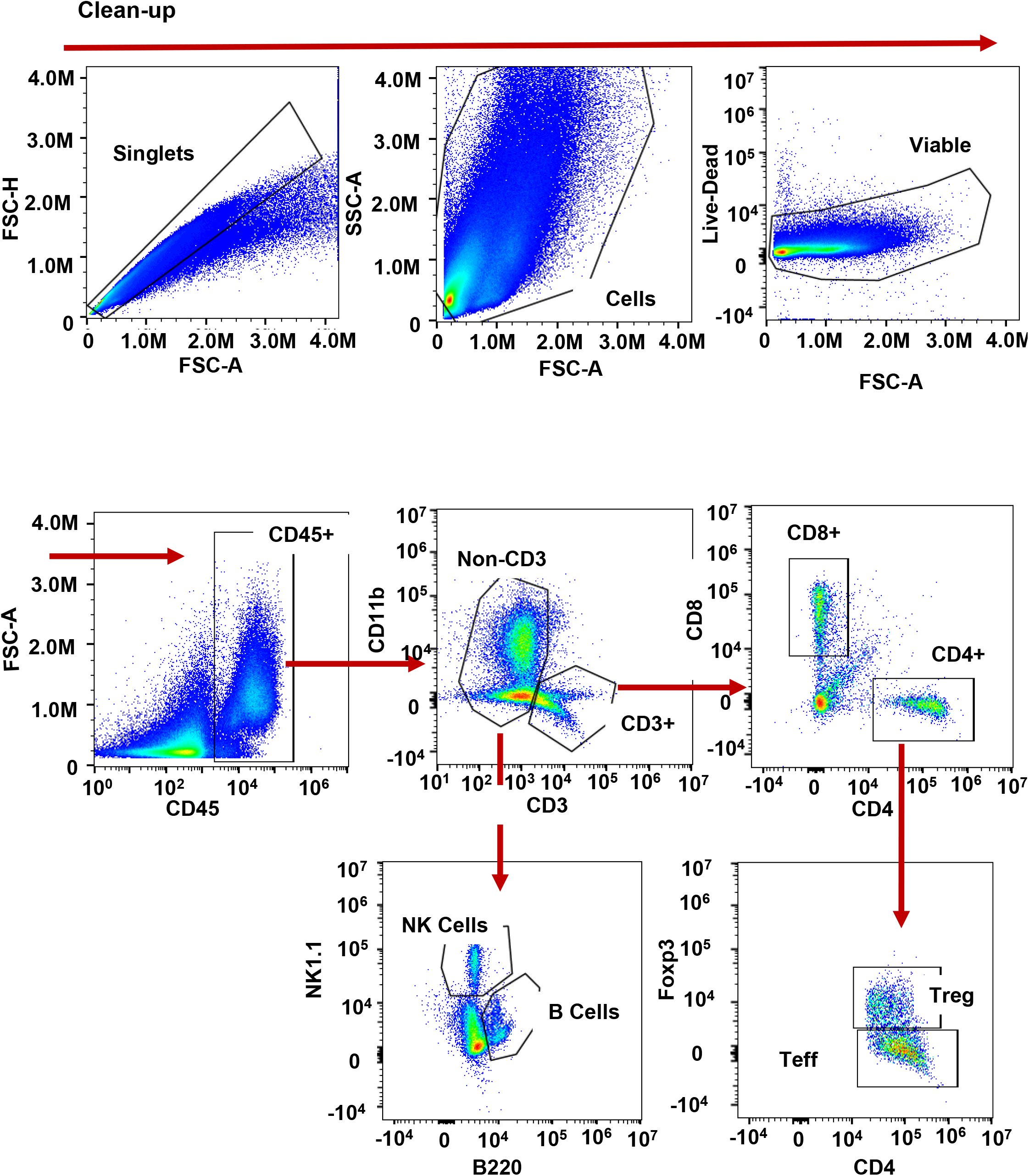
Representation of flow cytometry gating.

**Supplementary Figure 10.**
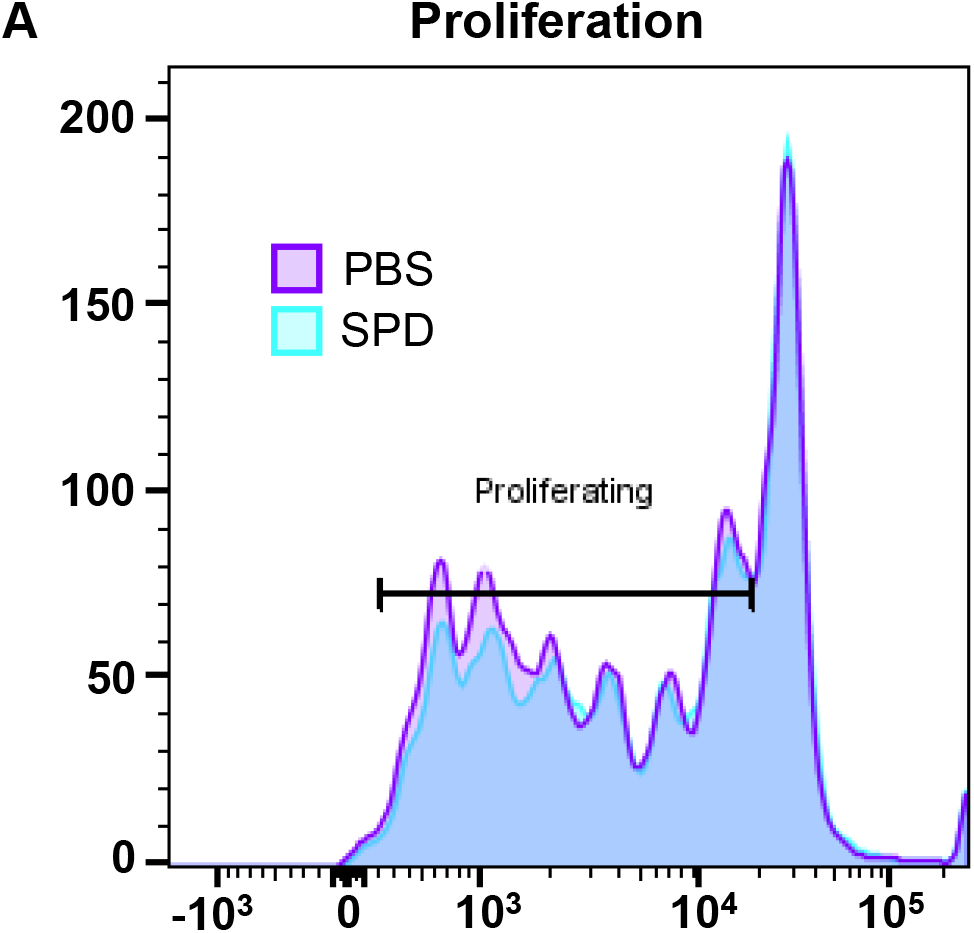
CD8+ T cells do not exhibit proliferation changes in the presence of SPD. Splenocyte-derived CD8+ T cells were treated with SPD or PBS vehicle *in vitro*. (A) CD8+ cells stained with CellTrace Violet and analyzed with flow cytometry; percentage of proliferating cells indicated by gate.

**Supplementary Figure 11.**
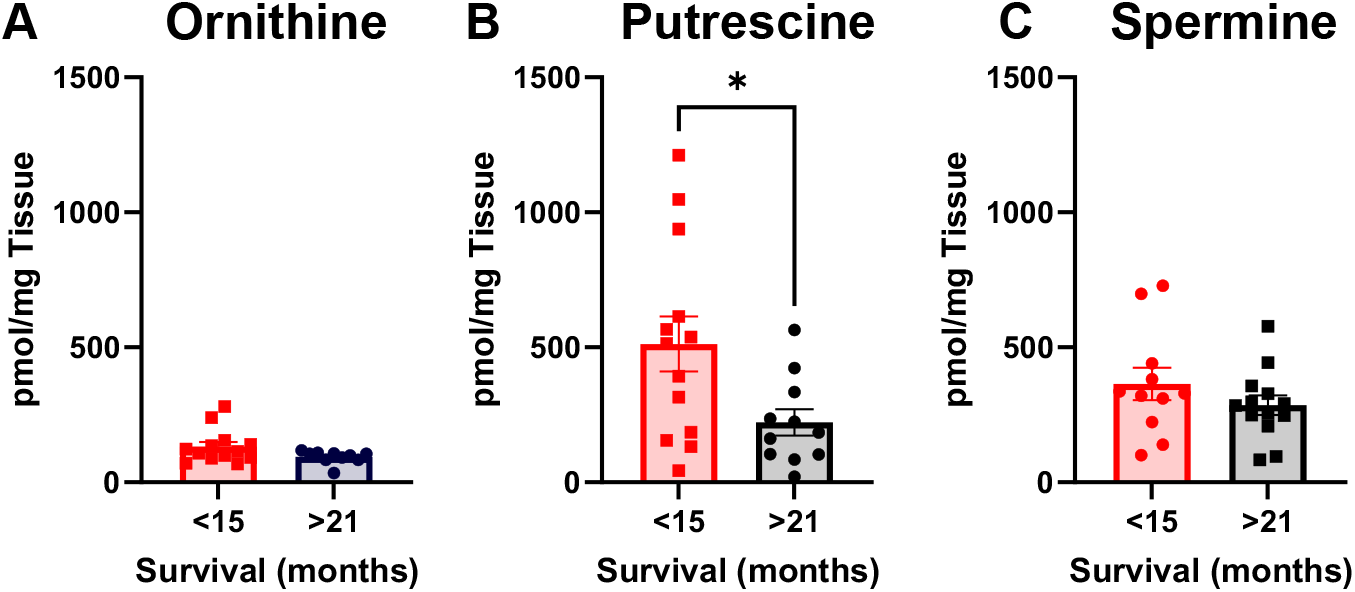
Additional members of polyamine pathway measured in GBM patient tumors. (A-C) Tumor tissue from primary resection of age-matched patients with longer survival compared to shorter survival; metabolites measured via LC-MS/MS. Statistical significance was determined by unpaired *t-*test (\**p*<0.05).

## Notes

### Competing Interest Statement

The authors have declared no competing interest.

## References Cited

1. Stupp, R. et al. High-grade glioma: ESMO Clinical Practice Guidelines for diagnosis, treatment and follow-up. Ann Oncol 25 Suppl 3, iii93-101 (2014).

2. Bell, E. H. et al. Molecular-Based Recursive Partitioning Analysis Model for Glioblastoma in the Temozolomide Era: A Correlative Analysis Based on NRG Oncology RTOG 0525. JAMA Oncol 3, 784– 792 (2017).

3. Furnari, F. B. et al. Malignant astrocytic glioma: genetics, biology, and paths to treatment. Genes Dev 21, 2683–710 (2007).

4. Ries, C. H. et al. Targeting tumor-associated macrophages with anti-CSF-1R antibody reveals a strategy for cancer therapy. Cancer Cell 25, 846–59 (2014).

5. Chaput, N. et al. Baseline gut microbiota predicts clinical response and colitis in metastatic melanoma patients treated with ipilimumab. Ann Oncol 28, 1368–1379 (2017).

6. Frankel, A. E. et al. Metagenomic Shotgun Sequencing and Unbiased Metabolomic Profiling Identify Specific Human Gut Microbiota and Metabolites Associated with Immune Checkpoint Therapy Efficacy in Melanoma Patients. Neoplasia 19, 848–855 (2017).

7. Fecci, P. E. et al. Increased regulatory T-cell fraction amidst a diminished CD4 compartment explains cellular immune defects in patients with malignant glioma. Cancer Res 66, 3294–302 (2006).

8. Jacobs, J. F. et al. Regulatory T cells and the PD-L1/PD-1 pathway mediate immune suppression in malignant human brain tumors. Neuro Oncol 11, 394–402 (2009).

9. Lewis, C. E. & Pollard, J. W. Distinct role of macrophages in different tumor microenvironments. Cancer Res 66, 605–12 (2006).

10. Platten, M., Wick, W. & Weller, M. Malignant glioma biology: role for TGF-beta in growth, motility, angiogenesis, and immune escape. Microsc Res Tech 52, 401–10 (2001).

11. Bayik, D. et al. Myeloid-Derived Suppressor Cell Subsets Drive Glioblastoma Growth in a Sex-Specific Manner. Cancer Discov. 10, 1210–1225 (2020).

12. Chongsathidkiet, P. et al. Sequestration of T cells in bone marrow in the setting of glioblastoma and other intracranial tumors. Nat Med 24, 1459–1468 (2018).

13. Watson, D. C. et al. GAP43-dependent mitochondria transfer from astrocytes enhances glioblastoma tumorigenicity. *Nat*. Cancer 4, 648–664 (2023).

14. Bayik, D. et al. Distinct Cell Adhesion Signature Defines Glioblastoma Myeloid-Derived Suppressor Cell Subsets. Cancer Res. 82, 4274–4287 (2022).

15. Rhun, E. L. et al. Molecular targeted therapy of glioblastoma. Cancer Treat. Rev. 80, (2019).

16. Shakya, S. et al. Altered lipid metabolism marks glioblastoma stem and non-stem cells in separate tumor niches. Acta Neuropathol. Commun. 9, 101 (2021).

17. Kant, S. et al. Enhanced fatty acid oxidation provides glioblastoma cells metabolic plasticity to accommodate to its dynamic nutrient microenvironment. Cell Death Dis. 11, 1–13 (2020).

18. Di Ianni, N., Musio, S. & Pellegatta, S. Altered Metabolism in Glioblastoma: Myeloid-Derived Suppressor Cell (MDSC) Fitness and Tumor-Infiltrating Lymphocyte (TIL) Dysfunction. Int. J. Mol. Sci. 22, 4460 (2021).

19. Hernández, A., Domènech, M., Muñoz-Mármol, A. M., Carrato, C. & Balana, C. Glioblastoma: Relationship between Metabolism and Immunosuppressive Microenvironment. Cells 10, 3529 (2021).

20. Pegg, A. E. Mammalian polyamine metabolism and function. IUBMB Life 61, 880–94 (2009).

21. Nowotarski, S. L., Woster, P. M. & Casero, R. A. Polyamines and cancer: implications for chemotherapy and chemoprevention. Expert Rev Mol Med 15, e3 (2013).

22. Miska, J. et al. Polyamines drive myeloid cell survival by buffering intracellular pH to promote immunosuppression in glioblastoma. Sci. Adv. 7, eabc8929 (2021).

23. Tangella, A. V., Gajre, A. S., Chirumamilla, P. C. & Rathhan, P. V. Difluoromethylornithine (DFMO) and Neuroblastoma: A Review. Cureus 15, e37680.

24. Khan, A. et al. Dual targeting of polyamine synthesis and uptake in diffuse intrinsic pontine gliomas. Nat. Commun. 12, 971 (2021).

25. Moulinoux, J. P., Quemener, V., Le Calve, M., Chatel, M. & Darcel, F. Polyamines in human brain tumors. A correlative study between tumor, cerebrospinal fluid and red blood cell free polyamine levels. J Neurooncol 2, 153–8 (1984).

26. Lee, J. et al. Sex-Biased T-cell Exhaustion Drives Differential Immune Responses in Glioblastoma. Cancer Discov. 13, 2090–2105 (2023).

27. Tavelin, B. & Malmström, A. Sex Differences in Glioblastoma—Findings from the Swedish National Quality Registry for Primary Brain Tumors between 1999–2018. J. Clin. Med. 11, 486 (2022).

28. Ostrom, Q. T., Rubin, J. B., Lathia, J. D., Berens, M. E. & Barnholtz-Sloan, J. S. Females have the survival advantage in glioblastoma. Neuro-Oncol. 20, 576–577 (2018).

29. Orrego, E. et al. Distribution of tumor-infiltrating immune cells in glioblastoma. CNS Oncol. 7, CNS21 (2018).

30. Han, S. et al. Tumour-infiltrating CD4+ and CD8+ lymphocytes as predictors of clinical outcome in glioma. Br. J. Cancer 110, 2560–2568 (2014).

31. Puleston, D. J. et al. Polyamine metabolism is a central determinant of helper T cell lineage fidelity. Cell 184, 4186–4202.e20 (2021).

32. Mandal, S., Mandal, A. & Park, M. H. Depletion of the polyamines spermidine and spermine by overexpression of spermidine/spermine N1-acetyltransferase 1 (SAT1) leads to mitochondria-mediated apoptosis in mammalian cells. Biochem. J. 468, 435–447 (2015).

33. Ruiz-Moreno, C. et al. Harmonized single-cell landscape, intercellular crosstalk and tumor architecture of glioblastoma. 2022.08.27.505439 Preprint at 10.1101/2022.08.27.505439 (2022).

34. Ravi, V. M. et al. Spatially resolved multi-omics deciphers bidirectional tumor-host interdependence in glioblastoma. Cancer Cell 40, 639–655.e13 (2022).

35. Alban, T. J. et al. Glioblastoma Myeloid-Derived Suppressor Cell Subsets Express Differential Macrophage Migration Inhibitory Factor Receptor Profiles That Can Be Targeted to Reduce Immune Suppression. Front. Immunol. 11, 1191 (2020).

36. Hibino, S., et al. Tumor cell–derived spermidine is an oncometabolite that suppresses TCR clustering for intratumoral CD8+ T cell activation. Proc. Natl. Acad. Sci. 120, e2305245120 (2023).

37. Yuan, H., Wu, S.-X., Zhou, Y.-F. & Peng, F. Spermidine Inhibits Joints Inflammation and Macrophage Activation in Mice with Collagen-Induced Arthritis. J. Inflamm. Res. 14, 2713–2721 (2021).

38. Zhou, S. et al. Reprogramming systemic and local immune function to empower immunotherapy against glioblastoma. Nat. Commun. 14, 435 (2023).

39. Guo, Y. et al. Spermine synthase and MYC cooperate to maintain colorectal cancer cell survival by repressing Bim expression. Nat. Commun. 11, 3243 (2020).

40. Peng, Q. et al. The Emerging Clinical Role of Spermine in Prostate Cancer. Int. J. Mol. Sci. 22, 4382 (2021).

41. Prasher, P. et al. Spermidine as a promising anticancer agent: Recent advances and newer insights on its molecular mechanisms. Front. Chem. 11, (2023).

42. Pietrocola, F. et al. Spermidine reduces cancer-related mortality in humans. Autophagy 15, 362–365 (2018).

43. Akinyele, O. & Wallace, H. M. Characterising the Response of Human Breast Cancer Cells to Polyamine Modulation. Biomolecules 11, 743 (2021).

44. Prados, M. D. et al. Phase III trial of accelerated hyperfractionation with or without difluromethylornithine (DFMO) versus standard fractionated radiotherapy with or without DFMO for newly diagnosed patients with glioblastoma multiforme. *Int*. J. Radiat. Oncol. 49, 71–77 (2001).

45. Liu, R. et al. Spermidine endows macrophages anti-inflammatory properties by inducing mitochondrial superoxide-dependent AMPK activation, Hif-1α upregulation and autophagy. Free Radic. Biol. Med. 161, 339–350 (2020).

46. Hu, C. et al. Polyamines from myeloid-derived suppressor cells promote Th17 polarization and disease progression. Mol. Ther. 31, 569–584 (2023).

47. Puleston, D. J. et al. Polyamines and eIF5A Hypusination Modulate Mitochondrial Respiration and Macrophage Activation. Cell Metab. 30, 352–363.e8 (2019).

48. Dono, A. et al. Glioma and the gut–brain axis: opportunities and future perspectives. Neuro-Oncol. Adv. 4, vdac054 (2022).

49. Patrizz, A. et al. Glioma and temozolomide induced alterations in gut microbiome. Sci. Rep. 10, 21002 (2020).

50. Raskov, H., Orhan, A., Christensen, J. P. & Gögenur, I. Cytotoxic CD8+ T cells in cancer and cancer immunotherapy. Br. J. Cancer 124, 359–367 (2021).

51. Lee, J., Kay, K., Troike, K., Ahluwalia, M. S. & Lathia, J. D. Sex Differences in Glioblastoma Immunotherapy Response. Neuromolecular Med. 24, 50–55 (2022).

52. Schildge, S., Bohrer, C., Beck, K. & Schachtrup, C. Isolation and culture of mouse cortical astrocytes. J. Vis. Exp. JoVE 50079 (2013) doi:10.3791/50079.

53. Analysis of polyamines as carbamoyl derivatives in urine and serum by liquid chromatography– tandem mass spectrometry - Byun - 2008 - Biomedical Chromatography - Wiley Online Library. https://analyticalsciencejournals.onlinelibrary.wiley.com/doi/10.1002/bmc.898.

